# Computational Analysis of Energy Landscapes Reveals Dynamic Features that Contribute to Binding of Inhibitors to CFTR-Associated Ligand

**DOI:** 10.1101/720342

**Authors:** Graham T. Holt, Jonathan D. Jou, Nicholas P. Gill, Anna U. Lowegard, Jeffrey W. Martin, Dean R. Madden, Bruce R. Donald

## Abstract

PDZ domains are small protein-binding domains that interact with short, mostly C-terminal peptides and play important roles in cellular signaling and the trafficking and localization of ion channels. The CFTR-associated ligand PDZ domain (CALP) binds to the cystic fibrosis transmembrane conductance regulator (CFTR) and mediates degradation of mature CFTR through lysosomal pathways. Inhibition of the CALP:CFTR interaction has been explored as a potential therapeutic avenue for cystic fibrosis (CF).^1^ Previously, we reported^2^ the ensemble-based computational design of a novel 6-residue peptide inhibitor of CALP, which resulted in the most binding-efficient inhibitor of CALP to date. This inhibitor, kCAL01, was designed using OSPREY^3^ and displayed significant biological activity in *in vitro* cell-based assays. Here, we report a crystal structure of kCAL01 bound to CALP (PDB ID: 6OV7). To elucidate the structural basis for the enhanced binding efficiency of kCAL01, we compare this structure to that of a previously developed inhibitor of CALP, iCAL36 (PDB ID: 4E34). In addition to performing traditional structural analysis, we compute the side-chain energy landscapes for each structure using the recently developed *MARK\** partition function approximation algorithm.^4^ Analysis of these energy landscapes not only enables approximation of binding thermodynamics for these structural models of CALP:inhibitor binding, but also foregrounds important structural features and reveals dynamic features, both of which contribute to the comparatively efficient binding of kCAL01. The investigation of energy landscapes complements traditional analysis of the few low-energy conformations found in crystal structures, and provides information about the entire conformational ensemble that is accessible to a protein structure model. Finally, we compare the previously reported NMR-based design model ensemble for kCAL01 vs. the new crystal structure and show that, despite the notable differences between the CALP NMR model and crystal structure, many significant features are successfully captured in the design ensemble. This suggests not only that ensemble-based design captured thermodynamically significant features observed *in vitro*, but also that a design algorithm eschewing ensembles would likely miss the kCAL01 sequence entirely.

**Graphical TOC Entry:** 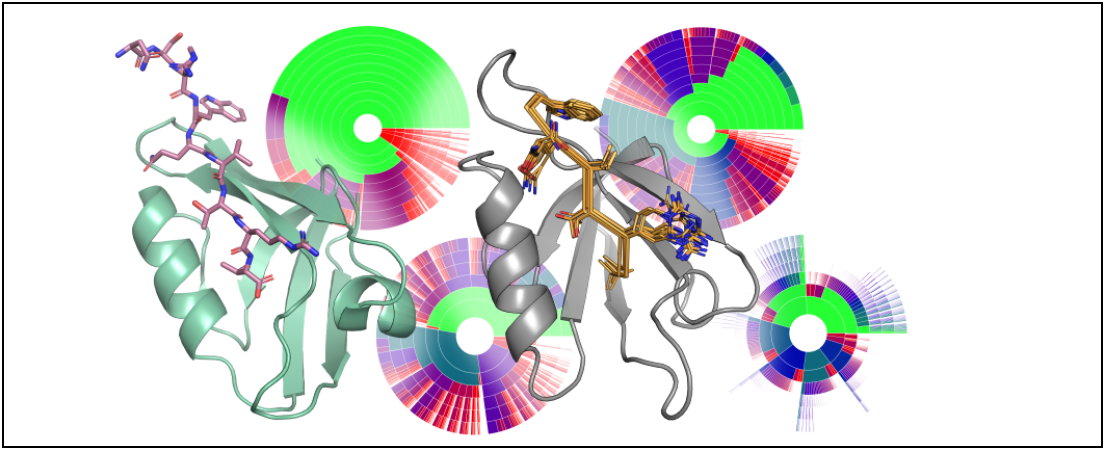

## 1 Introduction

Interactions between proteins and short linear motif peptides are important in many cellular contexts.^5^ One such class of peptide-binding proteins is the PDZ (PSD-95, discs large, ZO-1) domain family, characterized by an 80-90 residue motif^6^ that adopts a conserved fold comprised of 2-3 *α*-helices and 5-6 *β*-strands and binds C-terminal peptides through *β*-sheet interactions.^7^ These domains commonly modulate protein localization and complex assembly^8–10^ and regulate cell signalling,^11,12^ thereby playing critical roles in auditory and visual systems;^13,14^ epilepsy, pain, and addiction;^10,15^ synapse formation;^9^ cancer;^11,12,16,17^ and cystic fibrosis.^18–21^

Protein-peptide interactions have been implicated as therapeutic targets in cystic fibrosis (CF), a genetic disease characterized by defects in the cystic fibrosis transmembrane conductance regulator (CFTR) that result in impaired chloride ion transport.^18^ Approximately 90% of CF patients are homozygous or heterozygous for the *F508del (c.1521_1523delCTT)* mutation,^18,22,23^ which encodes a protein variant F508del-CFTR (p.Phe508del) with severe loss of function. In particular, CFTR is recycled from the cell membrane, and interactions of the CFTR C-terminus with the CFTR-associated ligand PDZ domain (CALP) preferentially target it for lysosomal degradation.^19,20^ Conversely, knockdown of CALP has been shown to rescue transepithelial chloride transport.^24^ Hence, inhibition of the interaction between the CFTR C-terminal peptide and CALP is a potential therapeutic avenue for CF. Understanding of the CALP:CFTR binding interaction is critical for the development of therapeutic inhibitors.

Previous work toward inhibitor development^1,25–27^ resulted in extensive characterization of the structural and stereochemical components of CALP binding. The structure of CALP bound to the CFTR C-terminal peptide was solved by solution NMR^25^ with well-resolved interactions between the 4 C-terminal peptide residues (P^-3^-P^0^) and CALP. This structure revealed canonical class 1 PDZ interactions^28^ including those between Leu P^0^ and a hydrophobic pocket between secondary structure elements *α*2 and *β*2, and the highly conserved hydrogen bond between Thr P^-2^ and His341. Peptide screening and iterative optimization using substitutional analysis^26^ revealed significant affinity effects of residues at other peptide positions up to P^-9^ and resulted in a decapeptide inhibitor (iCAL36) with an affinity of 22.6 *±* 8.0 *µ*M^26,27^ that rescued functional CFTR activity as assessed by *in vitro* Ussing chamber assays.^29^ Crystal structures of iCAL36 (and substituted peptide variants) in complex with CALP^1,27^ revealed structural features that influence CALP binding and selectivity. In particular, shifts in peptide orientation and location, along with conformational shifts in the carboxylate-binding loop (characterized by a *X*Φ_1_*G*Φ_2_ sequence motif, where Φ_*i*_ represents a hydrophobic amino acid), affect the binding geometry and specificity of the peptide P^0^ residue,^27^ allowing CALP to accommodate both Leu and Ile at P^0^. Additionally, side chain interactions at P^-1^, P^-3^, P^-4^, and P^-5^ modulate affinity and specificity of CALP binding.^1^ Finally, despite the fact that CALP:CFTR binding is thought to be primarily driven by enthalpic effects,^30^ NMR data and Molecular Dynamics (MD) simulations suggest that entropy may play a role in modulating CALP binding,^25^ a hypothesis which is reflected in studies of other PDZ domains.^31–34^

Previously,^2^ we developed the most binding efficient^35^ inhibitor of CALP to date using the OSPREY^3^ protein design software package, suggesting that components of CALP binding can be effectively captured using provable, ensemble-based computational protein design algorithms. Starting from the solution NMR structure of CALP:CFTR,^25^ we used the *K\** algorithm^36^ to compute approximations to *K*_*a*_ – the *K\** score – for CALP binding to *≈*8000 hexameric C-terminal peptides (residue positions P^-5^-P^0^). Retrospective predictions on 6223 previously characterized sequences^26,30^ showed that our algorithm was able to effectively classify sequences by binding affinity to CALP, with an area under the receiver operating characteristic (ROC) curve of 0.84. Additionally, OSPREY designs on 2166 sequences resulted in novel peptides that bind tightly to CALP. All of the top 11 prospective predictions were experimentally shown to bind with high affinity to CALP, and the tightest binding hexamer, kCAL01 (Ac-WQVTRV), bound with *K*_*i*_ = 2.3 *±* 0.2 *µ*M.^2^ kCAL01 bound more tightly than both the previous best hexamer inhibitor (iCAL35, WQT-SII, *K*_*i*_ = 14.0 *±* 1 *µ*M)^2^ and the best unmodified decamer (iCAL36, ANSRWPTSII, *K*_*i*_ = 22.6 *±* 8.0 *µ*M).^27^ Despite its small size (MW 829), kCAL01 binds with affinity comparable to a much larger (MW 1502) fluorescein-modified version of iCAL36 (*F\**-iCAL36, *K*_*d*_ = 1.3 *±* 0.1 *µ*M),^26^ yielding a much better binding efficiency for kCAL01 (See Table S1). Furthermore, kCAL01 rescued chloride ion transport activity of F508del-CFTR in cell-based assays.^2,37^ Ensemble-based design algorithms were shown to be critical for the success of this design: Ranking by energy of the global minimum energy conformations (GMECs) resulted in poor prediction accuracy and little overlap with the ensemble-based predictions.^2^ These data not only suggest that computational structural protein design (CSPD) algorithms can capture features that contribute to CALP:peptide binding, but also that ensemble-based or entropic effects are critical for prediction accuracy.

Indeed, computational designs are more biophysically accurate when they model protein thermodynamic ensembles.^2,36,38–45^ The objectives of CSPD algorithms are to 1) compute biophysical or thermodynamic properties of a protein or protein complex and 2) efficiently search for optimal sequences given an objective function. Without loss of generality, we choose binding affinity as our biophysical property of interest. CSPD algorithms search over a user-specified *input model* (viz., a structural model, allowed side chain and backbone flexibility, allowed mutations, energy function, etc.^3^). Because proteins exist as thermodynamic ensembles,^39,46^ principled algorithms should exploit statistical thermodynamics of non-covalent binding, and therefore require approximation of the partition function.^39,47^ However, because the conformation space available to proteins *in vivo* and *in vitro* is massive and grows exponentially with the number of flexible amino-acid residues, protein design algorithms often make simplifying modelling assumptions to allow tractable computation. Such assumptions often include 1) modeling only rigid, discrete side chain configurations, or *rotamers*,^48^ and a small set of discrete backbone conformations,^49–52^ 2) considering (or approximating) only a single global minimum energy conformation (GMEC),^50,53–58^ and 3) approximating the partition function using stochastic, heuristic sampling methods.^49,59,60^ However, these assumptions 1) fail to model small, commonly observed side chain and backbone movements, 2) entirely omit conformational entropy, and 3) often fail to find even the GMEC.^52^ Algorithms in the OSPREY software package efficiently solve protein design problems without these simplifications.^3^

Recently we developed the *MARK\** algorithm, which, in addition to provably and efficiently approximating partition functions for input model states (i.e. bound complex, unbound protein and ligand), allows visualization of the entire energy landscape accessible to a protein input model.^4^ *MARK\** provably bounds the energy and statistical weight of every conformation in the input model conformation space, allowing designers to compute and visualize changes in *conformation distribution*, instead of merely analyzing changes to a small set of low-energy conformations, or to ensemble averaged values like free energy or *K*_*a*_. By computing both a conformation distribution and provably approximating energies for every conformation, *MARK\** enables visualization of the energy landscape. This novel capability complements traditional structural analysis by providing insight into entropic and dynamic contributions to binding.

In this work, we report a 1.7 Å resolution crystal structure of a decapeptide variant of the peptide inhibitor kCAL01 bound to CALP (PDB ID: 6OV7). To evaluate the structural basis for the enhanced binding efficiency of kCAL01, we compare this structure to that of a previously developed decapeptide inhibitor of CALP, iCAL36 (PDB ID: 4E34).^27^ In addition to performing traditional structural analysis, we compute energy landscapes for bound and unbound structural models for CALP:kCAL01 and CALP:iCAL36 using *MARK\**. From these landscapes we compute approximations to the free energy, internal energy, and entropy for each model state, and use these quantities to model thermodynamics of binding for CALP:kCAL01 and CALP:iCAL36. Additionally, we analyze the energy landscapes for dynamic effects, and show that these energy landscapes foreground important structural features of binding and reveal dynamic features that may contribute to the efficient binding of kCAL01. We conclude that investigation of energy landscapes complements traditional analysis of one or few low-energy structures represented in crystal structures, and provides important information about the entire conformational ensemble that is available to a protein structure model. Finally, to assess the extent to which designs reported in Ref.^2^ are a result of accurate modeling of structural and dynamic components of CALP binding, we compare the design output ensemble for kCAL01 to the newly solved crystal structure. We show that, despite notable differences between the NMR-based CALP structure used as design input vs. the bound crystallographic conformation of CALP, many significant crystal structure features are captured in the design output ensemble. This suggests that the success of the ensemble-based computational design of kCAL01^2^ was a result of effective modeling of structural and dynamic features of binding.

## 2 Methods

### 2.1 Structure Determination of CALP:kCAL01

Recombinant CAL PDZ (CALP; UniProt accession number Q9HD26-2; residues 278-362) was expressed and purified as described previously.^61^ Briefly, an expression construct was engineered in pET16b containing a N-terminal decahistidine tag, a short linker, and a human rhinovirus (HRV) 3C protease cleavage site, followed by the CALP sequence. The construct was transformed into E. *coli* BL21 (DE3) RIL cells, expression was induced as previously described^61^ (except that TB medium was used), and protein was purified by nickel-nitrilotriacetic acid (NiNTA) affinity chromatography and size-exclusion chromatography (SEC). Following removal of the affinity tag and linker by HRV-3C protease cleavage, CALP was recovered in the flow-through fraction of a NiNTA affinity column and further purified by SEC. To facilitate crystallization, kCAL01 was synthesized as a decapeptide (*ANSR*WQVTRV) containing four N-terminal residues (in italics) that form lattice contacts in other CALP:peptide co-crystals.^1,27,61,62^ The kCAL01 decapeptide was synthesized using standard Fmoc solid-phase peptide synthesis and purified by reverse-phase HPLC. Peptide mass was confirmed using liquid chromatography/mass spectrometry (LC/MS). Using the hanging-drop method, CALP:kCAL01 co-crystals were obtained by mixing 1 mM kCAL01 and 6 mg/mL CALP with reservoir solution containing 25% (*w/v*) PEG 8000, 5% (*v/v*) PEG 400, 150 mM sodium chloride, and 100 mM Tris pH 8.5. Crystals were transferred into cryoprotectant solution (25% [*w/v*] PEG 8000, 15% [*v/v*] PEG 400, 150 mM sodium chloride, and 100 mM Tris pH 8.5) and flash-cooled in a liquid-nitrogen bath. Oscillation diffraction data were recorded at 100 K on beamline BL9-2 at the Stanford Synchrotron Radiation Lightsource (SSRL) over a 180° range with 0.5 s, 0.2° exposures. Reflection intensities were integrated and scaled using the XDS package^63^ (version 20190315). Initial phase estimates were obtained by molecular replacement using Phaser^64^ within the Phenix package^65^ (version 1.15.2) and using PDB ID 4E34^27^ as the search model (containing chains A and C only). Subsequent model building and refinement were performed using Phenix and Coot^66^ (version 0.8.9.2) to generate the final model of CALP in complex with the kCAL01 decapeptide at 1.71 Å resolution. Data quality and refinement statistics are reported in Table 1. The coordinates and structure factors have been deposited in the Protein Data Bank (www.rcsb.org) with ID 6OV7.

**Table 1:** Data collection and refinement statistics for CALP:kCAL01 complex (PDB ID: 6OV7).

|  |  |
| --- | --- |
| **Data collection** |  |
| Space group | $P 2_1 2_1 2_1$ |
| Unit cell dimensions $a, b, c$ (Å) | 43.6, 60.8, 81.2 |
| Resolution<sup>a</sup> (Å) | 48.6 – 1.71 (1.83 – 1.71) |
| $R_{\text{sym}}^b$ (%) | 10.8 (95.9) |
| $I/\sigma_I$ | 14.0 (2.0) |
| Completeness (%) | 99.6 (99.8) |
| **Refinement** |  |
| Total number of reflections | 23681 |
| Reflections in test set | 1160 |
| $R_{\text{work}}^c / R_{\text{free}}^d$ (%) | 18.2 / 21.4 (28.1 / 30.7) |
| Number of atoms<sup>e</sup> |  |
| Protein | 1470 |
| Water | 175 |
| Ramachandran plot<sup>f</sup> (%) | 98.4, 1.6, 0, 0 |
| $B_{\text{av}}$ (Å<sup>2</sup>) | |
| Protein | 22.3 |
| Solvent | 33.1 |
| Bond length RMSD (Å) | 0.01 |
| Bond angle RMSD (°) | 1.18 |

<sup>a</sup>Values in parentheses are for data in the highest resolution shell. <sup>b</sup> $R_{\text{sym}} = \sum_h \sum_i |I(h) - I_i(h)| / \sum_h \sum_i I_i(h)$ , where $I_i(h)$ and $I(h)$ values are the $i$ -th and mean measurements of the intensity of reflection $h$ .
<sup>c</sup> $R_{\text{work}} = \sum ||F_{\text{obs}}|_h - |F_{\text{calc}}|_h| / \sum |F_{\text{obs}}|_h$ , $h \in \{\text{working set}\}$ . <sup>d</sup> $R_{\text{free}}$ is calculated as $R_{\text{work}}$ for the reflections $h \in \{\text{test set}\}$ . <sup>e</sup>Includes alternate conformations. <sup>f</sup>Core, allowed, generously allowed, disallowed.

### 2.2 Computational Methods

The new crystal structure of CALP:kCAL01 (PDB ID: 6OV7, protomer A) and the crystal structure of CALP:iCAL36 (PDB ID: 4E34, protomer A) were used to model energy landscapes of CALP binding to kCAL01 and iCAL36, respectively. For each structure, we computed energy landscapes for three states: the bound CALP:peptide complex, the unbound CALP, and the unbound peptide. This was accomplished by first defining a set of accessible conformations for each state, and then using the *MARK\** algorithm^4^ in OSPREY 3.0^3^ to compute both a provable approximation to the partition function value and an approximation to the energy landscape.

Sets of accessible conformations, or *conformation spaces*, were defined as follows for each state. These conformation spaces are an approximation to the ensemble of conformations available to each state *in vivo*. First, hydrogens were added to each crystal structure using the MolProbity server^67^ in order to generate *protonated crystal structures*. Backbone atom coordinates for the bound complex state were obtained directly from protonated crystal structures 6OV7 (for the CALP:kCAL01 state) and 4E34 (for the CALP:iCAL36 state). Nine residues for CALP and the six most C-terminal residues for kCAL01 or iCAL36 (for a total of 15 residues in in each bound complex, see Table S2) were modeled as continuously flexible using continuous rotamers^68,69^ in OSPREY. As in Refs.,^4,44,68,70^ rotamers from the Penultimate Rotamer Library^48^ were allowed to adopt any side-chain conformation such that all *χ*-angles are within *±*9° of their modal *χ*-angles. For all other residues, side-chain coordinates were obtained from protonated crystal structures. Models for unbound CALP and peptide states were obtained by removing all atoms of the peptide or CALP structure, respectively, from the complex state. Thus, we defined approximations to the conformational ensembles for bound and unbound states, herein referred to as *models*, for CALP:kCAL01 and CALP:iCAL36.

For each model, we computed *ε*-approximate bounds on the value of the partition function to a deterministic, guaranteed accuracy of *ε <* 0.01 using the *MARK\** algorithm^4^ in OSPREY. All computations were run on 40-48 core Intel Xeon nodes with up to 500 GB of memory. As proved previously,^4^ not only does *MARK\** compute a provable *ε*-approximation to the partition function, it also bounds the energy landscape by provably approximating the energy and therefore statistical weight of all model conformations in the conformation space.

### 2.3 Entropy, Internal Energy, and Helmholtz Free Energy Calculation

Aggregate values for the ensembles in each state were computed by bounding the energy for each conformation in the ensemble, and combining these energy bounds. For each bound and unbound state, we first computed bounds on the the energy of each conformation in the conformational ensemble defined by that state, as was done in Ref.,^2–4,42,71,72^ described in Section 2.2. Using these energy bounds, we then computed bounds on corresponding Boltzmann-weighted partition function 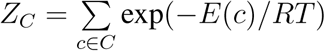, where *E* is a function that returns the energy of conformation *c*, by computing and summing bounds on the Boltzmann weights for conformations *c* in that state *C* (See Ref.^4^ for details). We then divided the upper bound on Boltzmann weight of *c* by the upper bound on the partition function to compute the probability *p*_*c*_ for each conformation *c* within the ensemble. Using these probabilities, we then calculated the entropy 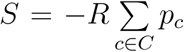 ln *p*_*c*_ and internal energy 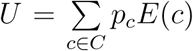 of the ensemble, and combined these two values to compute the Helmholtz free energy *F* = *U −T S* at a temperature of 298.15 K. Here, *E* is a function that returns the lower bound on the energy of conformation *c*. We also used the upper bounds on Boltzmann weight for each conformation to compute energy landscape diagrams,^4^ which are explained in Section 2.4.

Although the *K\** score exhibits good Spearman’s rank correlation with experimental *K*_*a*_ values,^3,71^ the correlation between *K\** scores and *K*_*a*_ is not yet quantitative. First, most physics-based energy functions are based on small-molecule energetics, which can overestimate van der Waals terms and thereby overestimate internal energy. Additionally, the input models used in current computation model only a subset of biologically available flexibility – in this case, flexibility was restricted to up to 4 side-chain *χ* angles per residue. We did not model explicit waters, instead relying on the EEF1 implicit solvation model^73^ in osprey. As a result, our models likely underestimate entropy and overestimate internal energy. Therefore, we scaled our thermodynamic values, decreasing internal energy *U* by a factor of 4, similar to the method described in Ref.^4^

### 2.4 Interpretation of energy landscape diagrams

Visual representations of computed energy landscapes can be found in Figures 4 and 5. For a full description of ring diagram visualizations, see Ref.^4^ Briefly, each concentric ring represents a design amino acid residue, with each ring arc representing a single rotamer assignment to that residue given the residue assignments of the inner arcs, or “partial conformation.” Therefore, any arc in the outermost ring represents a “full conformation”, where all amino acid positions are each assigned a single rotamer. The angle of any given arc corresponds to the partition function contribution of all conformations that contain the given partial conformation. The color of any given arc corresponds to the smallest energy difference between the GMEC and the lowest energy conformation that contains the given partial conformation, with small energy differences colored green, and larger energy differences colored red. Notably, white gaps are indicative of relatively high energy conformations that individually contribute less than 0.1% of the partition function value. Therefore, a ring diagram visually represents the entire energy landscape for a design problem, showing the distribution of conformations according to their probability.

**Figure 1:**
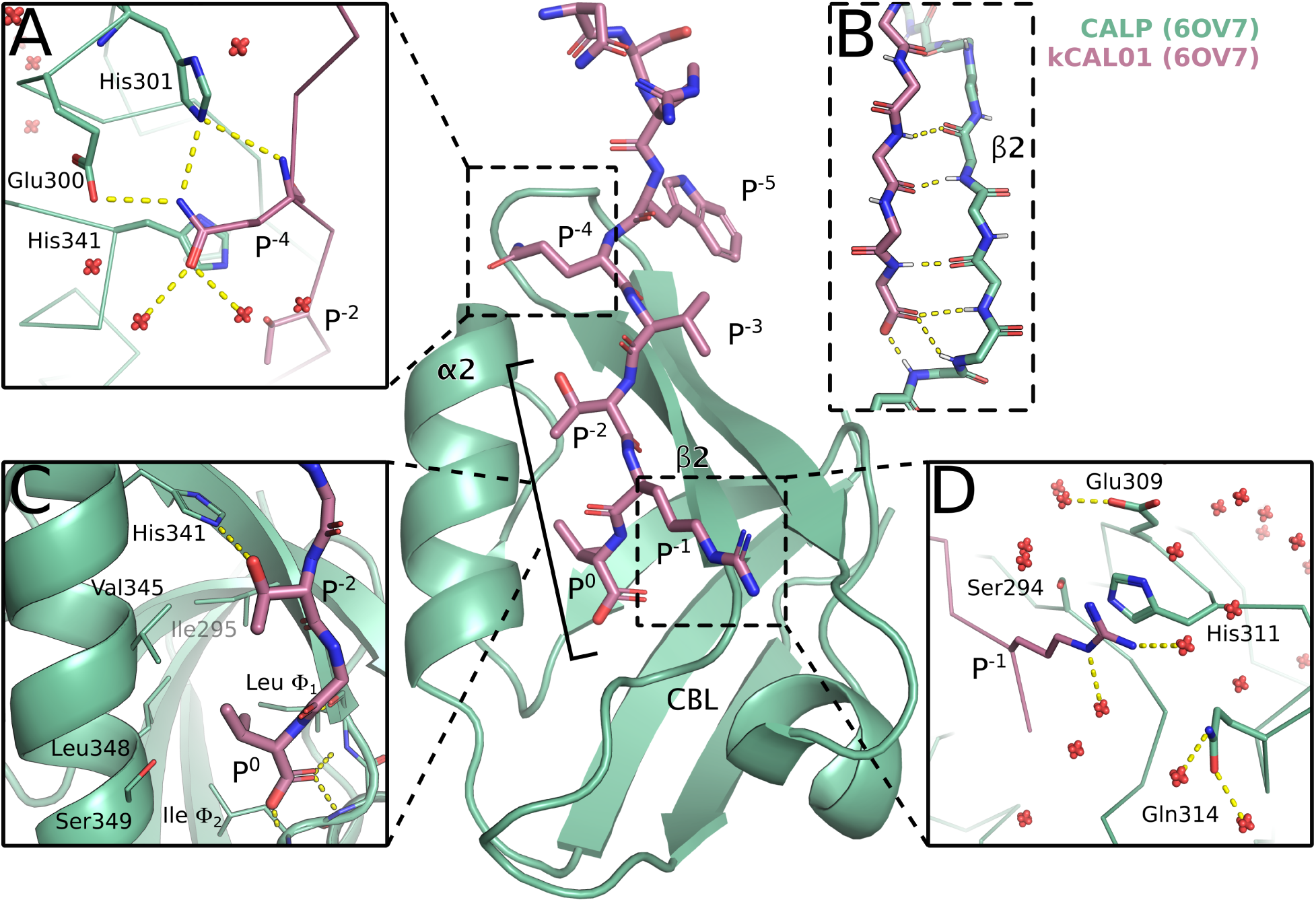
Crystal structure of CALP:kCAL01 (PDB ID: 6OV7) displays canonical class 1 PDZ binding and favorable interactions at P^-1^ and P^-4^. The CALP and kCAL01 crystal structure is shown in green and pink, respectively (protomer A). Hydrogen bonds predicted by the Probe^75^ software are represented as dashed yellow lines. The kCAL01 peptide binds in the groove defined by helix *α*2 and strand *β*2. (A) Gln P^-4^ makes favorable hydrogen bonds with Glu300 or His301, and forms van der Waals interactions with His341. Additionally, Gln P^-4^ can coordinate with several water molecules, shown as red, non-bonded oxygens. (B) kCAL01 binds in the groove defined by helix *α*2 and strand *β*2 and forms an anti-parallel *β*-sheet interaction with strand *β*2. The C-terminus of kCAL01 forms favorable hydrogen bonds with the carboxylate-binding loop (CBL). (C) kCAL01 displays features of class 1 PDZ binding,^28^ including the conserved hydrogen bond between Thr P^-2^ and His341, as well as the interaction between the Val P^0^ side chain and the hydrophobic pocket. (D) Arg P^-1^ appears to form favorable *π*-interactions with His311, and also coordinates with several waters.

**Figure 2:**
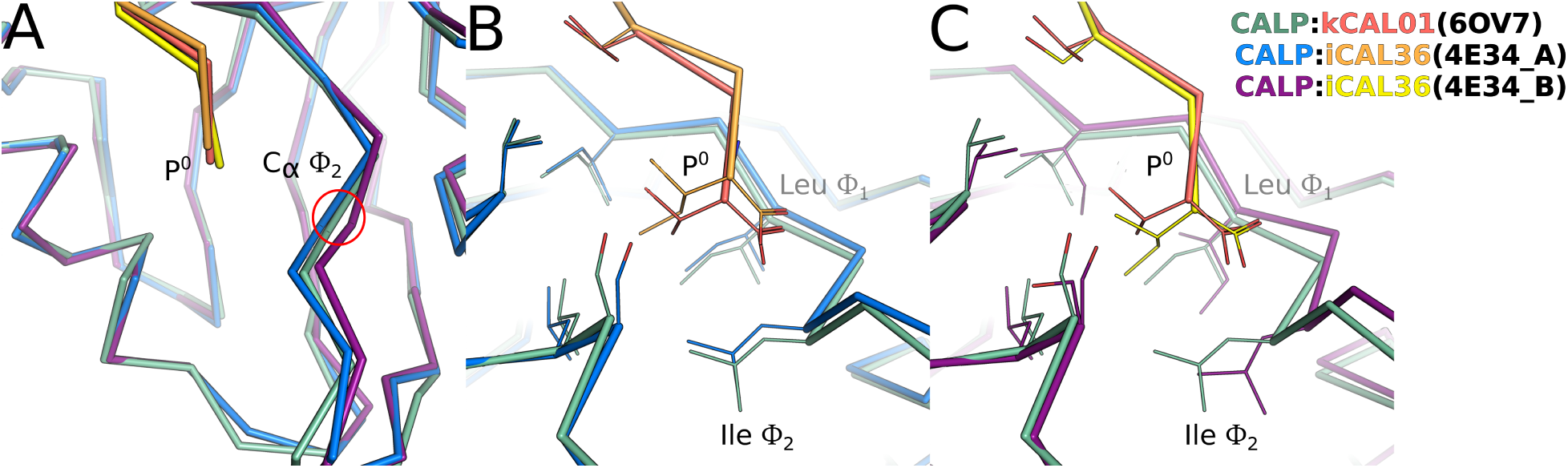
Binding geometry of kCAL01 P^0^ and the carboxylate-binding loop. Superimposed views of the P^0^ interaction with the carboxylate-binding loop (CBL) and hydrophobic pocket (side chains shown as lines) are shown for CALP:kCAL01 protomer A (green:pink), CALP:iCAL36 protomer A (blue:orange), and CALP:iCAL36 protomer B (purple:yellow). (A) Superimposed C_*α*_ traces show that the CALP conformation at the Ile Φ_2_ C_*α*_ is more similar to the CALP:iCAL36 protomer A conformation than the CALP:iCAL36 protomer B conformation. (B) A pairwise comparison shows that the CALP:kCAL01 CBL geometry matches most closely with CALP:iCAL36 protomer A, seen at side chains at CBL positions Φ_1_ and Φ_2_. However, the kCAL01 peptide P^0^ shifts toward the CBL by 0.7 Å relative to the CALP:iCAL36 structure. (C) A pairwise comparison shows that the CALP:kCAL01 peptide orientation matches most closely with CALP:iCAL36 protomer B, seen at position P^0^. However, the CALP:iCAL36 CBL shifts outward by 1.3 Å relative to the CALP:kCAL01 structure, and the hydrophobic pocket expands due to changes in rotamer at CBL position Φ_1_.

**Figure 3:**
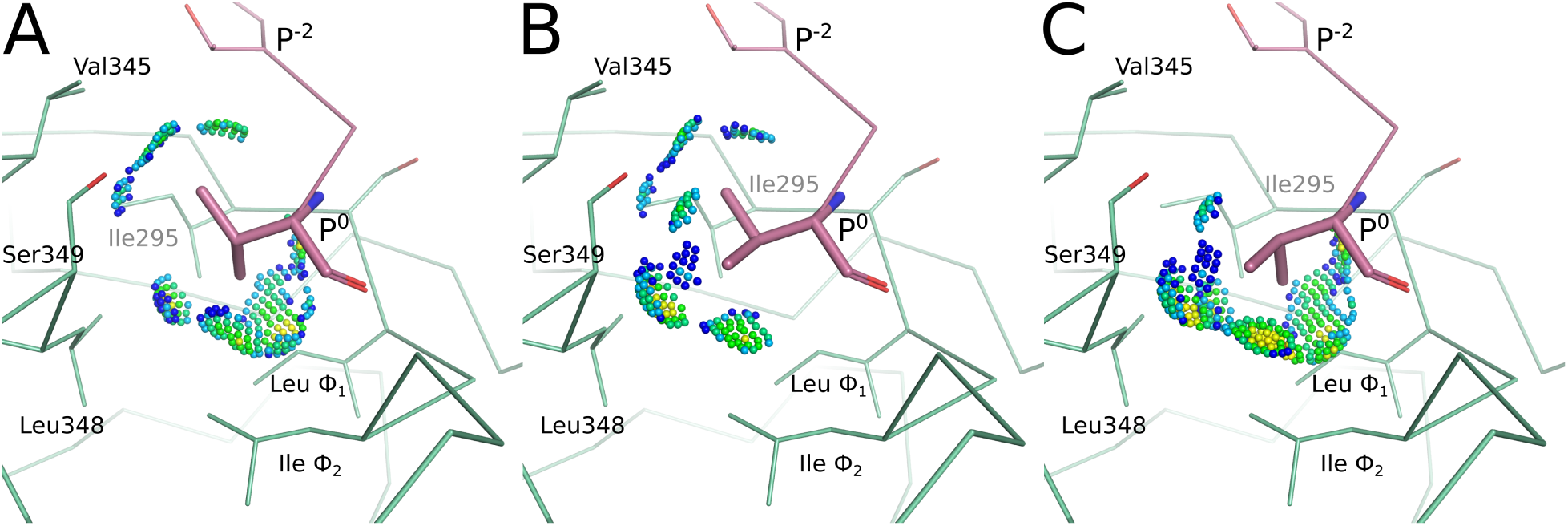
Energy landscape analysis reveals conformational heterogeneity at Val P^0^ for CALP:kCAL01. Energy landscape analysis of bound kCAL01 indicates three rotamers at peptide P^0^ that contribute significantly to the partition function. We refer to these rotamers as **t, p**, or **m**, which describe the valine N-C_*α*_-C_*β*_ -C_*γ*1_ dihedral angle as *trans* (∼ 180°), *plus 60* (∼ 60°), or *minus 60* (*∼ −*60°), respectively, conforming to the convention defined in Ref.^48^ Conformations containing each of these three rotamers were selected from the bound kCAL01 ensemble. Interactions between the P^0^ side-chain atoms and the CALP structure are shown using Probe dots,^75^ where green and blue dots indicate favorable interactions, yellow dots indicate small overlaps, and red and pink lines show steric clashes. All three of these rotamers form favorable interactions with the hydrophobic binding pocket, and our analysis suggests that the complex can sample any of these rotamers with relatively high probability. (A) The **p** rotamer forms favorable interactions with Thr P^-2^, Val345, Leu348, Ile Φ_2_, and Leu Φ_1_. (B) The **t** rotamer forms favorable interactions with Thr P^-2^, Val345, Ser349, Leu348, and Ile Φ_2_. The slight overlaps (yellow dots) generated due to the interaction with Leu348, along with the lack of interaction with Leu Φ_1_ suggest that this conformation is slightly less favorable than the **p** rotamer. (C) The **m** rotamer forms favorable interactions with Ser349, Leu348, Ile Φ_2_, and Leu Φ_1_. Slight overlaps (yellow dots) can be seen in interactions with Leu348, Ile Φ_2_, and Leu Φ_1_, and there is no interaction with Val345, suggesting that this rotamer may be slightly less favorable than either the **p** or **t** rotamers. Nevertheless all three rotamers are well sampled in the ensemble, and contribute significantly to the partition function.

**Figure 4:**
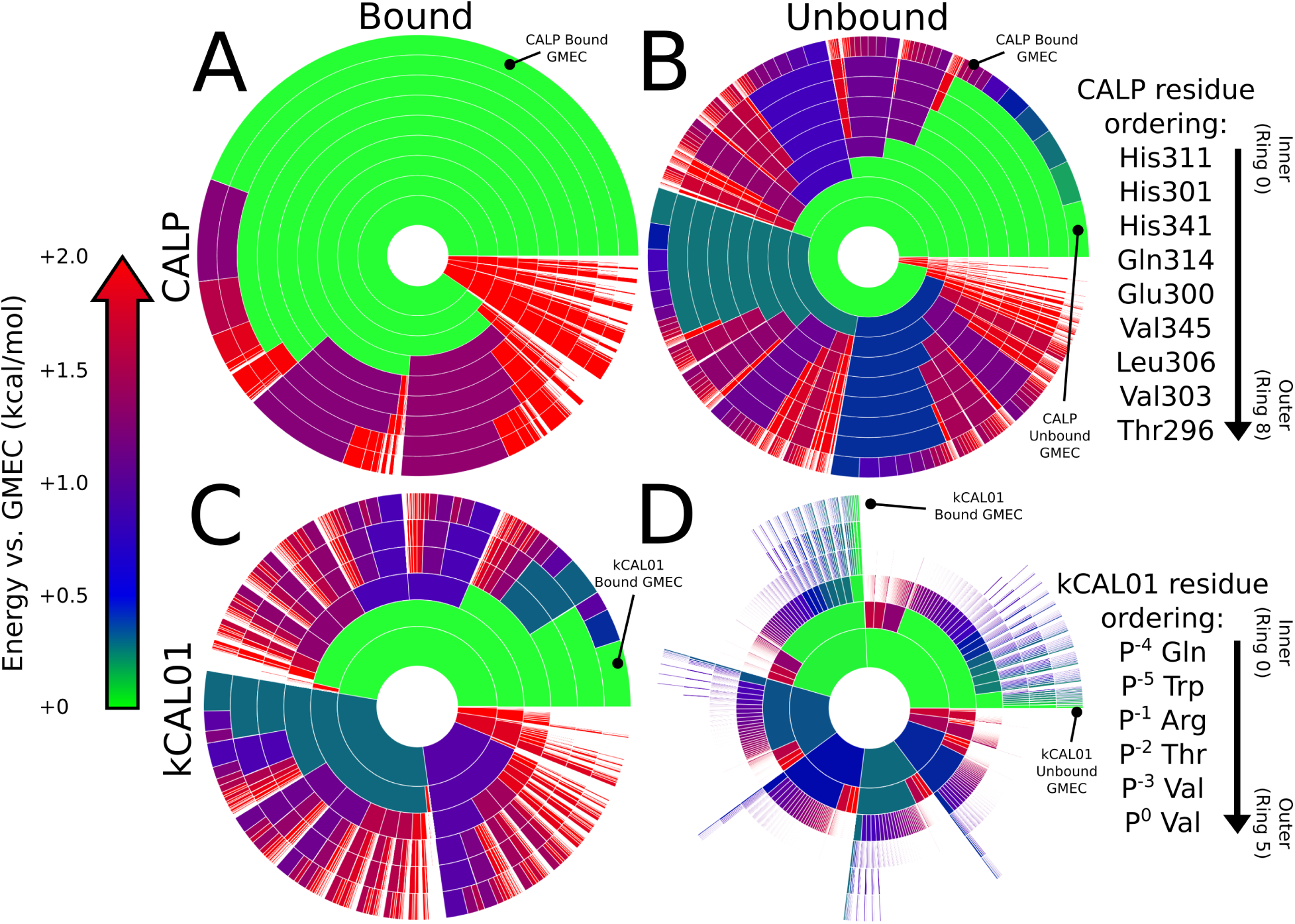
Energy landscape analysis reveals components of binding thermodynamics for CALP:kCAL01. Upper bounds on the Boltzmann-weighted partition function computed using the *MARK\** algorithm^4^ in OSPREY^3^ for a 15-residue design at the protein-protein interface of CALP:kCAL01 are shown as colored ring charts. A brief explanation of the ring chart diagram can be found in Section 2.4. (A, B) Energy landscapes for CALP in the bound (A) and unbound (B) states show the change in conformation distribution induced by binding. (B) Unbound CALP shows a wide conformation distribution, with the unbound GMEC accounting for roughly 5% of the partition function, with conformational entropy generated largely by residues Thr296 and His301, indicated by a large number of similarly-sized arcs at their corresponding rings. (A) In contrast, bound CALP has a narrow distribution, with the GMEC accounting for nearly 50% of the partition function. Importantly, the bound and unbound GMECs are not the same conformation, and do not have the same energy. (C, D) Energy landscapes for kCAL01 in the bound (C) and unbound (D) states show a similar change in conformation distribution upon binding. (D) Unbound kCAL01 shows a very high entropy conformation distribution, with many conformations that contribute to the partition function, as seen by the presence of many small arcs in the outer ring. This is consistent with our expectations for an extended peptide backbone. (C) Conversely, the bound kCAL01 energy landscape shows that the GMEC accounts for roughly 5% of the partition function. Even so, the landscape suggests considerable entropy, driven by residues at P^-4^, P^-1^, and P^-2^, and P^0^. Prediction of conformational heterogeneity at P^0^ is particularly interesting given its buried location. Thermodynamic parameters calculated from these partition functions (See Sec. 3.2.2) indicate that binding of kCAL01 to CALP results in a decrease in internal energy and a decrease in entropy, which is represented visually in these energy landscapes.

**Figure 5:**
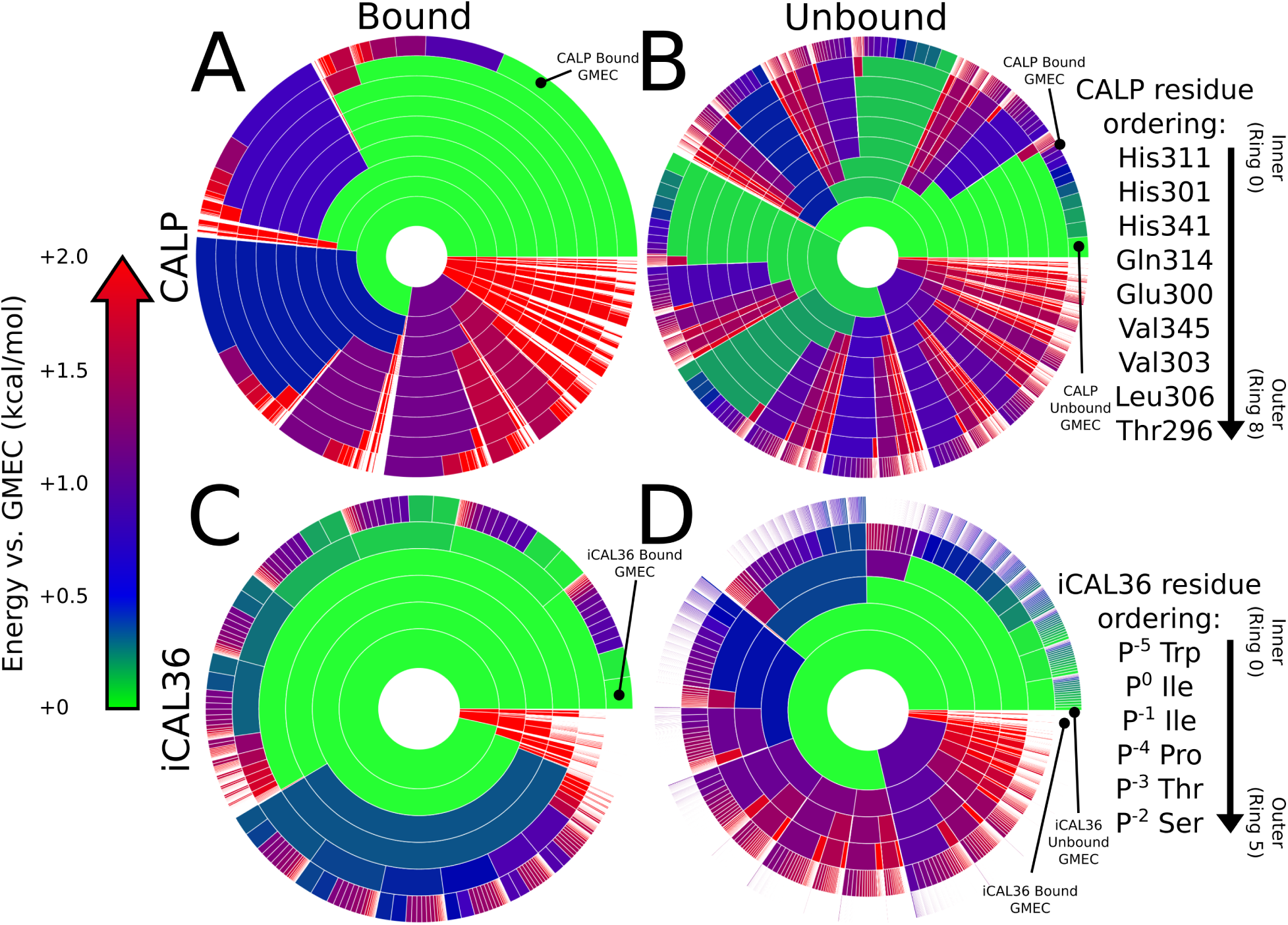
Energy landscape analysis reveals components of binding thermodynamics for CALP:iCAL36. Upper bounds on the Boltzmann-weighted partition function computed using the *MARK\** algorithm^4^ in OSPREY^3^ for a 15-residue design at the protein-protein interface of CALP:iCAL36 are shown as colored ring charts. A brief explanation of the ring chart diagram can be found in Section 2.4. (A, B) Energy landscapes for CALP in the bound and unbound states show the change in conformation distribution induced by binding. (B) Unbound CALP shows a wide conformation distribution, with the unbound GMEC accounting for roughly 2% of the partition function, with conformational variation in multiple residues, indicated by a large number of similarly-sized arcs at multiple rings. (A) Bound CALP has a narrower distribution, with the GMEC accounting for roughly 20% of the partition function. (C,D) Energy landscapes for iCAL36 in the bound and unbound states show a change in conformation distribution upon binding. (D) Unbound iCAL36 has a high entropy conformation distribution, with many conformations that contribute to the partition function, as seen by the presence of many small arcs in the outer ring. This is consistent with our expectations for an extended peptide backbone. (C) The bound iCAL36 energy landscape exhibits a lower entropy distribution, with the GMEC accounting for roughly 5% of the partition function. Even so, the landscape suggests considerable heterogeneity, much of which is attributable to variation at P^-2^, and P^-3^. Interestingly, in contrast to the bound kCAL01 landscape, little heterogeneity is predicted at P^0^. Thermodynamic parameters calculated from these partition functions (See Sec. 3.2.2) indicate that binding of iCAL36 to CALP results in a decrease in internal energy and a decrease in entropy, which is represented visually in these energy landscapes.

## 3 Results and Discussion

### 3.1 Structural analysis of CALP:kCAL01

Co-crystals were formed with recombinant CALP and a decapeptide variant of kCAL01 with a four-residue N-terminal extension (*ANSR*WQVTRV, extension in italics). The refined model exhibits excellent fit to the density, with *R*_work_ and *R*_free_ of 0.182 and 0.214, respectively, and lies within typical peptide geometry constraints (Table 1). The asymmetric unit consists of two protomers (A and B) of CALP complexed with kCAL01 (Figure S1A). The 9 or 8 C-terminal residues of the peptide ligand are well-resolved in protomers A and B, respectively. The crystal structure was deposited as PDB ID: 6OV7.

Alignment of protomers A and B of 6OV7 by CALP main-chain atoms using PyMOL^74^ results in good overlap, with 281 of the 348 total backbone atoms aligning with an RMSD of 0.32 Å. Notable differences can be seen between the two protomer structures at two sites: helix *α*1 and adjacent to the carboxylate-binding loop (CBL). Significant distortion of the protomer B *α*1 helix results from an inter-protomer disulfide bond between CALP residue 319 (protomer A) and 319 (protomer B) (Figure S1 A,B). We hypothesize that this disulfide bond and resulting helix *α*1 distortion are artifacts of crystallization. Pronounced conformational differences adjacent to the protomer A and B CBLs, which connect the *β*-strands *β*1 and *β*2, are evident between CALP residues 284 and 289 (Figure S1B) However, these differences occur upstream of the CBL sequence motif residues 291 (Φ_1_) and 293 (Φ_2_), and do not appear to affect peptide binding. Due to the distortion of protomer B helix *α*1, the following analysis focuses on the protomer A CALP:kCAL01 structure.

#### 3.1.1 Gross structural analysis of CALP:kCAL01 reveals canonical PDZ:peptide binding

The overall topology of the protomer A CALP fold, comprised of 5 *β*-strands and 2 *α*-helices, matches well with previous CALP structures and represents a canonical PDZ fold.^6,7^ Class 1 PDZ domains bind to peptides containing a C-terminal S/T-X-Φ binding motif, where Φ is a hydrophobic residue, and form an anti-parallel *β*-sheet with the *β*2 strand.^28^ kCAL01 binds in a manner consistent with typical class 1 PDZ domains, occupying the groove defined by helix *α*2 and strand *β*2 (Figure 1). Four main-chain hydrogen bonds are formed between CALP *β*2 and the 3 C-terminal kCAL01 residues P^-2^-P^0^, forming an anti-parallel *β*-strand interaction (Figure 1B). This positions the most C-terminal peptide residue (P^0^) such that the main-chain carboxyl terminus interacts with the CBL, defined by an *X*Φ_1_*G*Φ_2_ sequence motif, where Φ_*i*_ is a hydrophobic amino acid.^7,27^ Additionally, the hydrophobic P^0^ side chain is buried in the pocket defined by CALP residues Leu291, Ile293, Ile295, Val345, and Leu348 (Figure 1C). The CALP:kCAL01 structure also contains the critical hydrogen bond between Thr P^-2^ and His341, which plays a significant role in defining PDZ domain Class 1 specificity^28^ (Figure 1C). Notably, only the six C-terminal residues of the extended kCAL01 peptide form any direct contacts with CALP – the four additional N-terminal residues make only lattice contacts and were added to facilitate crystallization. Overall, this structure depicts a binding interaction that is consistent with the structural characteristics observed for canonical class 1 PDZ domains.^7,28^

To evaluate the basis for the enhanced efficiency of the CALP:kCAL01 binding interaction, we compared this crystal structure (PDB ID: 6OV7, protomer A) to the structure of CALP:iCAL36 (PDB ID: 4E34, protomers A and B), a previously developed inhibitor of CALP that also exhibits in-cell activity,^29^ but binds less tightly to CALP. We note that CALP residue numbering differs between these two structures, with numbering for 4E34 +8 relative to 6OV7. Unless otherwise noted, all residue numbering refers to the 6OV7 numbering convention.

#### 3.1.2 Comparison of CALP:kCAL01 to CALP:iCAL36 reveals differences in carboxylate-binding loop conformation

First, we analyzed the CBL conformation and peptide orientation, because previous work^27^ demonstrated that these features play a role in modulating CALP specificity for peptide residue P^0^. In particular, through analysis of 4E34, Ref.^27^ presented two structural mechanisms by which CALP accommodates a Ile P^0^ side chain: 1) a CBL conformation that narrows the entrance to the hydrophobic binding pocket concomitant with an N-terminal peptide shift (4E34, protomer A), and 2) a CBL conformation that widens the entrance to the hydrophobic binding pocket concomitant with a change in rotamer at Leu Φ_1_, thus expanding the hydrophobic binding pocket (4E34, protomer B). kCAL01, in contrast, has a valine at position P^0^. To investigate the structural consequences of this substitution, we aligned each protomer of CALP:iCAL36 to CALP:kCAL01 by the mainchain atoms of CALP secondary structure elements *β*2 and *α*2, which flank the peptide-binding groove. Structures of CALP:kCAL01 protomer A and CALP:iCAL36 protomer A showed good correspondence, with an RMSD of 0.24 Å, and CALP:kCAL01 protomer A and CALP:iCAL36 protomer B aligned with an RMSD of 0.41 Å.

The overall binding geometry of the CBL and peptide for CALP:kCAL01 contains distinct features of both CALP:iCAL36 protomers A and B (Figure 2). The CBL conformation of kCAL01 at residue Φ_2_ is more similar to that of the iCAL36 protomer A than protomer B, with C_*α*_ deviations of 0.5 Å and 1.3 Å respectively (See Figure 2A). Additionally, the rotamer at loop residue Φ_1_ matches with iCAL36 protomer A (Figure 2B). Overall, the CALP CBL when bound to kCAL01 adopts a conformation that narrows the entrance of the hydrophobic binding pocket, which is similar to the previously reported CALP:iCAL36 protomer A structure (Figure 2B). This suggests that kCAL01 binding to CALP does not require hydrophobic pocket expansion to accommodate the Val P^0^.

However, the bound kCAL01 peptide shifts toward the CBL, similar to the CALP:iCAL36 protomer B structure (Figure 2C). kCAL01 shifts toward the CBL relative to iCAL36 protomer A by 0.7 Å measured at the P^0^ C_*α*_ (Figure 2B). This results in side-chain positioning at kCAL01 Val P^0^ that is intermediate to CALP:iCAL36 protomers A and B. This shift propagates up the backbone of the peptide, as the P^-1^ C_*α*_ also shifts by 0.8 Å. These results suggest that the presence of a valine at P^0^, rather than a sterically larger leucine or isoleucine, allows shifts in the peptide backbone that accommodate the less common C-terminal side chain within a high-affinity interaction.

Based on this structural analysis, it is unclear how the changes in P^0^ binding mode between CALP:kCAL01 and CALP:iCAL36 affect binding affinity. On the one hand, the Val P^0^ present in kCAL01 appears to allow a peptide C-terminal shift without requiring a shift in the CBL and hydrophobic pocket expansion. On the other hand, it is not clear whether this C-terminal peptide shift is either favorable or unfavorable for binding. Indeed, the interactions formed by Val and Ile P^0^ in 6OV7 and 4E34, respectively, appear qualitatively similar, and the inclusion of the sterically larger Ile could more effectively fill the pocket. Overall, more analysis is needed to clarify the effects of structural variation at this site. This analysis is provided in Section 3.2.1, where investigation of CALP:kCAL01 and CALP:iCAL36 energy landscapes suggests that these structural shifts allow the kCAL01 Val P^0^ to sample three favorable rotamers, which we predict to be favorable for binding.

#### 3.1.3 Comparison of CALP:kCAL01 and CALP:iCAL36 at modulator residues reveals interactions that favor kCAL01 binding

Previous work^1,26,62^ pinpointed “modulator” residues at P^-1^, P^-3^, and P^-4^-P^-9^ that show individually modest effects on binding and specificity, but together can create significant effects. We compared the CALP:kCAL01 and CALP:iCAL36 protomer A structures in order to determine the effect of these modulator residues on inhibitor binding.

kCAL01 contains an arginine residue at P^-1^ that interacts with His311 in an apparent *π*-cation interaction (Figure 1D). In contrast, iCAL36 contains an isoleucine at this position, which forms minor van der Waals interactions with His311 and Ser294. These interactions appear much less extensive than those formed by the Arg P^-1^. Favorable interactions between CALP and kCAL01 are indicated at Gln P^-4^, which hydrogen bonds with Glu300 or His301, in addition to forming van der Waals interactions with His341 and interacting with several waters (Figure 1A). While the Pro P^-4^ found in the CALP:iCAL36 structure does make interactions with His301, His341, and a single water molecule, these interactions appear to be slightly less favorable.

kCAL01 and iCAL36 differ only slightly at P^-3^, containing a valine and threonine residue, respectively. Both residues form van der Waals interactions between a methyl group and Ser308, but only the kCAL01 Val P^-3^ forms interactions with Thr296. These differences, while minor, suggest slightly more favorable interactions for kCAL01 at this position.

Overall, the most notable structural differences in binding stereochemistry between kCAL01 and iCAL36 occur at residues P^-1^ and P^-4^, where mutations to long polar and charged residues likely result in an increase in favorable energetic interactions. These results suggest that sequence differences shift the thermodynamic balance: the more hydrophobic iCAL36 peptide may have higher energy alone in solvent, whereas the more polar kCAL01 sequence is preferentially stabilized in the bound state.

### 3.2 Energy landscape analysis of CALP:kCAL01

Conformational entropy can play a significant role in defining protein structure and function.^76–78^ For this reason, when modeling binding of protein:ligand complexes, it is useful to compute partition functions over protein ensembles to better model and understand binding thermodynamics.^2,36,38–44^ To approximate the conformational ensembles involved in CALP:kCAL01 and CALP:iCAL36 binding, we computed partition functions and energy landscapes using OSPREY for bound and unbound models of CALP:kCAL01 (PDB ID: 6OV7, protomer A) and CALP:iCAL36 (PDB ID: 4E34, protomer A) as described in Section 2.2. We compared energy landscape features of CALP:kCAL01 and CALP:iCAL36 to reveal dynamic features that contribute to CALP:kCAL01 binding. Furthermore, we used these energy landscapes to compute approximations to thermodynamic components of binding (described in Section 2.3) to analyze differences in binding of CALP:kCAL01 and CALP:iCAL36. A discussion on how to interpret energy landscape diagrams presented herein can be found in Section 2.4.

#### 3.2.1 Energy landscape comparison of CALP:kCAL01 and CALP:iCAL36 foregrounds structural and dynamic features that explain differences in binding

Detailed comparison of CALP:kCAL01 (Figure 4) and CALP:iCAL36 (Figure 5) landscapes reveals local differences in side-chain conformational distributions for each structure. This is most notable when comparing the bound inhibitor landscapes of kCAL01 and iCAL36 (Figures 4C, 5C, respectively). Note that for ease of comparison we have decomposed the bound complex CALP:kCAL01 landscape into bound CALP and kCAL01 landscapes. The original convolved landscapes can be found in Figure S3.

The bound kCAL01 landscape (Figure 4C) indicates that residue Val P^0^ adopts three significant rotamers, shown by subdivision into three arcs at the outermost ring (ring 5). In contrast, the bound iCAL36 landscape (Figure 5C) indicates that residue Ile P^0^ adopts only one significant rotamer, shown in the second ring from the center (ring 1). Structural analysis of the rotamer distribution for this residue (Figure 3) suggests that Val P^0^ forms favorable interactions with the CALP hydrophobic pocket in each of three rotamers defined by a rotation of ∼ 60° around the N-C_*α*_-C_*β*_-C_*γ*1_ dihedral angle. As a result, the landscape analysis of bound kCAL01 suggests that residue Val P^0^ interacts with low energy *and* locally high entropy. Conversely, the iCAL36 Ile P^0^ is likely too large to interact favorably in multiple rotameric states and is predicted to occupy only one significant rotamer. These predicted differences in conformational heterogeneity at P^0^ could help explain the improved binding efficiency of kCAL01. Such dynamic features are difficult to visualize when examining only a static crystal structure, but are now made clear by the energy landscape analysis.

Comparison of the energy landscapes of bound CALP for the CALP:kCAL01 and CALP:iCAL36 models (Figures 4A, 5A) also reveals differences in conformational heterogeneity. CALP bound to kCAL01 appears heavily conformationally restricted, with the GMEC occupying nearly 50% of the landscape (Figure 4A), while CALP bound to iCAL36 is less conformationally restricted, with the GMEC occupying roughly 20% of the landscape (Figure 5A). These differences appear to be driven in large part by differences in residue conformational heterogeneity at His311 and His301 (CALP:kCAL01 residue numbering, CALP:iCAL36 numbering is +8 relative), which can be seen by comparing the innermost two rings (ring 0 and 1) in the two bound CALP landscapes (Figures 4A, 5A). For each of ring 0 (the innermost) or 1 (the second innermost), the bound CALP landscape in the iCAL36 structure shows an additional minor rotamer population, shown by a purple or blue arc, respectively. This indicates that the rotamer distribution for His311 and His301 in the bound CALP:iCAL36 model has more entropy than that in the CALP:kCAL01 model.

Examining structural interactions between His311 and His301 and the peptide inhibitor for CALP:kCAL01 and CALP:iCAL36 structures suggests that this relative increase in entropy for CALP:iCAL36 can be explained by a loss of interactions between His311 and peptide P^-1^, and between His301 and peptide P^-4^. Specifically, kCAL01 Arg P^-1^ forms strong *π*-stacking interactions with His311, while iCAL36 Ile P^-1^ forms weaker interactions with the analogous His319. Similarly, kCAL01 Gln P^-4^ forms a hydrogen bond with His301, while iCAL36 Pro P^-4^ forms van der Waals interactions. As a result, we expect these histidines to form more energetically favorable interactions with kCAL01 than iCAL36. Our models predict that these favorable interactions are sensitive to the rotamer choice at His311 and His301, resulting in a less conformationally heterogeneous ensemble at these positions. Thus, energy landscape analysis both reveals conformational heterogeneity at Val P^0^ and draws attention to important modulator residue interactions in the bound state.

#### 3.2.2 Thermodynamics of CALP:kCAL01 and CALP:iCAL36 energy landscapes reveal decreases in internal energy and entropy upon binding

Energy landscapes of CALP:kCAL01 binding visualize the loss of entropy upon binding. Figure 4 depicts energy landscapes of the unbound CALP (panel B), unbound kCAL01 (panel D), and bound CALP and kCAL01 (panels A and C, respectively). Comparison of the unbound and bound ensemble landscapes for CALP (Figure 4A,B) reveals the significant loss of entropy due to conformational rearrangement upon binding. The unbound CALP landscape (Figure 4B) shows many low-energy conformations that contribute to the partition function, indicated by the many green-blue arcs in the outermost ring. Conversely, the bound ensemble of CALP (Figure 4A) is dominated by a single low-energy conformation, indicated by the large green arc, with the rest of the landscape occupied by higher-energy minor conformations. This indicates that the conformational rearrangement due to binding of kCAL01 imposes significant entropic cost that must be compensated for by the gain of favorable inter-molecular interactions.

Comparison of the unbound and bound ensemble landscapes for kCAL01 (Figure 4C and D, respectively) reveals a similar picture, with the decrease in entropy upon binding illustrated by the decrease in number and increase in size of the arcs in the outermost ring. Notably, for the unbound kCAL01 landscape, the GMEC contributes relatively little and outer rings are characterized by extensive white-space, indicating the presence of many conformations that occupy individually less than 0.1% of the partition function. Together, these features are indicative of a high entropy landscape. In contrast, the bound kCAL01 landscape is characterized by fewer conformations that contribute relatively more to the partition function, and a GMEC that occupies roughly 5% of the partition function. These landscape representations depict the loss of entropy upon binding of CALP:kCAL01.

Using these energy landscapes, we calculated approximations to the ensemble-weighted internal energy and entropy for the bound and unbound states of CALP:kCAL01 as described in Section 2.3. Additionally, we computed the same approximations for the binding-competent ensemble^79^ – an alchemical state defined by the conformations and occupancies found in the bound protein or ligand modeled with the energy field of the unbound state – to deconvolve the changes in entropy vs. internal energy upon binding. Conveniently, as shown previously^79^ this construction allows us to decompose binding into an “induced fit” step, involving a change in conformation distribution, and a “lock and key” step, in which protein:ligand interactions are formed, without regard to actual mechanism. At the chosen temperature of ∼298 K, both CALP and kCAL01 exhibit a change in conformation distribution upon binding. This change in distribution results in a change in internal energy and entropy for CALP of 0.113 kcal/mol and −1.97 kcal/mol, respectively, and for kCAL01 of 0.234 kcal/mol and −2.23 kcal/mol, respectively, quantifying the large decrease in entropy due to binding, with the total change in entropy due to binding *T*Δ*S*_binding_ of −4.19 kcal/mol. Complex formation – the “lock and key” step – results in a decrease in internal energy: whereas the combined internal energy of the unbound models is −18.84 kcal/mol, the bound model internal energy is −28.49 kcal/mol, resulting in a Δ*U*_binding_ of −9.65 kcal/mol. As a result, the approximated change in Helmholtz free energy due to binding Δ*F*_binding_ is −5.46 kcal/mol. These models suggest that both binding partners incur penalties to entropy and internal energy when adopting the binding competent ensemble, which are compensated for by a large decrease in internal energy upon complex formation.

Energy landscapes of CALP:iCAL36 binding reveal a loss of entropy upon binding that is similar to that of CALP:kCAL01. Figure 5 depicts energy landscapes of the unbound CALP (panel B), unbound iCAL36 (panel D), and bound CALP and iCAL36 (panels A and C, respectively). Comparison of bound and unbound states for CALP and iCAL36 also reveal a decrease in entropy upon binding for both binding partners, indicated by CALP and iCAL36 energy landscapes exhibiting a reduction in the number of arcs and an increase in arc size upon binding.

Approximations of ensemble-weighted internal energy and entropy for bound, binding competent ensemble, and unbound models of CALP:iCAL36 revealed a smaller decrease in entropy upon binding compared to CALP:kCAL01, but also showed a smaller decrease in internal energy upon binding. At a temperature of ∼298 K, our models indicate that both CALP and iCAL36 undergo a change in conformation distribution upon binding. This results in a change in internal energy and entropy for CALP of 0.004 kcal/mol and −1.60 kcal/mol, respectively, and for iCAL36 of 0.558 kcal/mol and −1.89 kcal/mol, respectively, illustrating a decrease in entropy due to binding, with the total change in entropy due to binding *T*Δ*S*_binding_ of −3.48 kcal/mol. Complex formation results in a decrease in internal energy: whereas the combined internal energy of the unbound models is −17.7 kcal/mol, the bound model internal energy is −25.6 kcal/mol, resulting in a Δ*U*_binding_ of −7.84 kcal/mol. As a result, the approximated change in Helmholtz free energy due to binding Δ*F*_binding_ is −4.36 kcal/mol. Similar to CALP:kCAL01, both binding partners undergo a loss of entropy and reduction in internal energy upon binding.

Overall, these landscapes and thermodynamic calculations suggest that although both CALP:kCAL01 and CALP:iCAL36 undergo a decrease in both entropy and internal energy upon binding, the energetic interactions gained upon binding are less favorable for CALP:iCAL36 than for CALP:kCAL01. This is reflected in the change in internal energy due to binding for each model, with CALP:kCAL01 and CALP:iCAL36 exhibiting Δ*U*_binding_ of −9.65 kcal/mol and Δ*U*_binding_ of −7.84 kcal/mol, respectively. Although the entropic penalty due to binding is less for CALP:iCAL36, these models predict that kCAL01 binds more tightly to CALP than does iCAL36, with Δ*F*_binding_ values of −5.46 kcal/mol and −4.36 kcal/mol, respectively. Despite the fact that our models account for only a subset of biologically relevant flexibility (See Section 2.3), this prediction is confirmed by experimental measurements^2,26,27^ (See Table S1), suggesting that these models are capturing biologically relevant features.

### 3.3 Design model corresponds closely with bound crystal structure

In this section, we briefly compare the design model reported in Ref.^2^ with 6OV7 to determine which key structural features contributed to a successful kCAL01 design. We perform structural comparisons for both the CALP:kCAL01 *design output model* (defined as the ensemble of structures output from the *K\** algorithm^36^ in OSPREY) and the CALP:CFTR *design input model* (defined as the structural input to OSPREY). A schematic diagram depicting these definitions is shown in Figure 6. We begin by comparing gross features, and then proceed to identify shared side-chain energetic and dynamic interactions.

**Figure 6:**
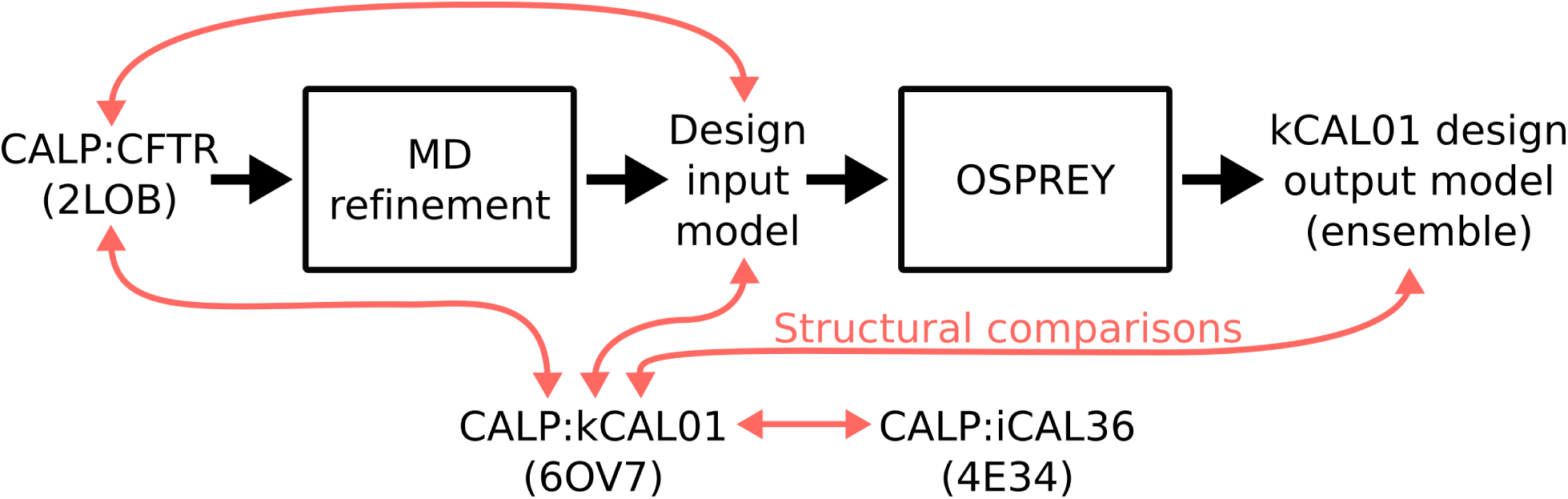
Schematic diagram of design process reported in Ref.^2^ and structural comparisons presented in this work. A flowchart of design work performed in Ref.^2^ is shown with black arrows. First, a design input model was generated by performing MD refinement of an NMR structure of CALP:CFTR.^25^ Using OSPREY, this input model was used for the *K\** algorithm’s^36^ ensemble-based design of peptide inhibitors of CALP, resulting in a design output model (ensemble) of CALP:kCAL01.^2^ Finally, in this work we perform detailed structural comparisons between several CALP:peptide structures and models, indicated by red arrows.

To determine the accuracy of the structural design model reported in Ref.,^2^ we compared the crystal structure of CALP:kCAL01 to the ensemble of 100 low energy structures that comprise the CALP:kCAL01 design output model^2^ (Figure 7). We aligned members of the design output ensemble to the CALP:kCAL01 crystal structure using the main chain of secondary-structure elements *α*2 and *β*2 and obtained good alignment quality, with one representative structure aligning with an RMSD of 0.94 Å. Deviations were primarily a result of a more relaxed hydrophobic binding pocket in the design model relative to the CALP:kCAL01 crystal structure 6OV7, involving an outward shift of the *β*2 strand (Figure 7A). Additionally the loop connecting *β*2 and *β*3 adopts a different conformation, resulting in a global loop shift and a significant change in orientation of the *β*2 strand (Figure 7A).

**Figure 7:**
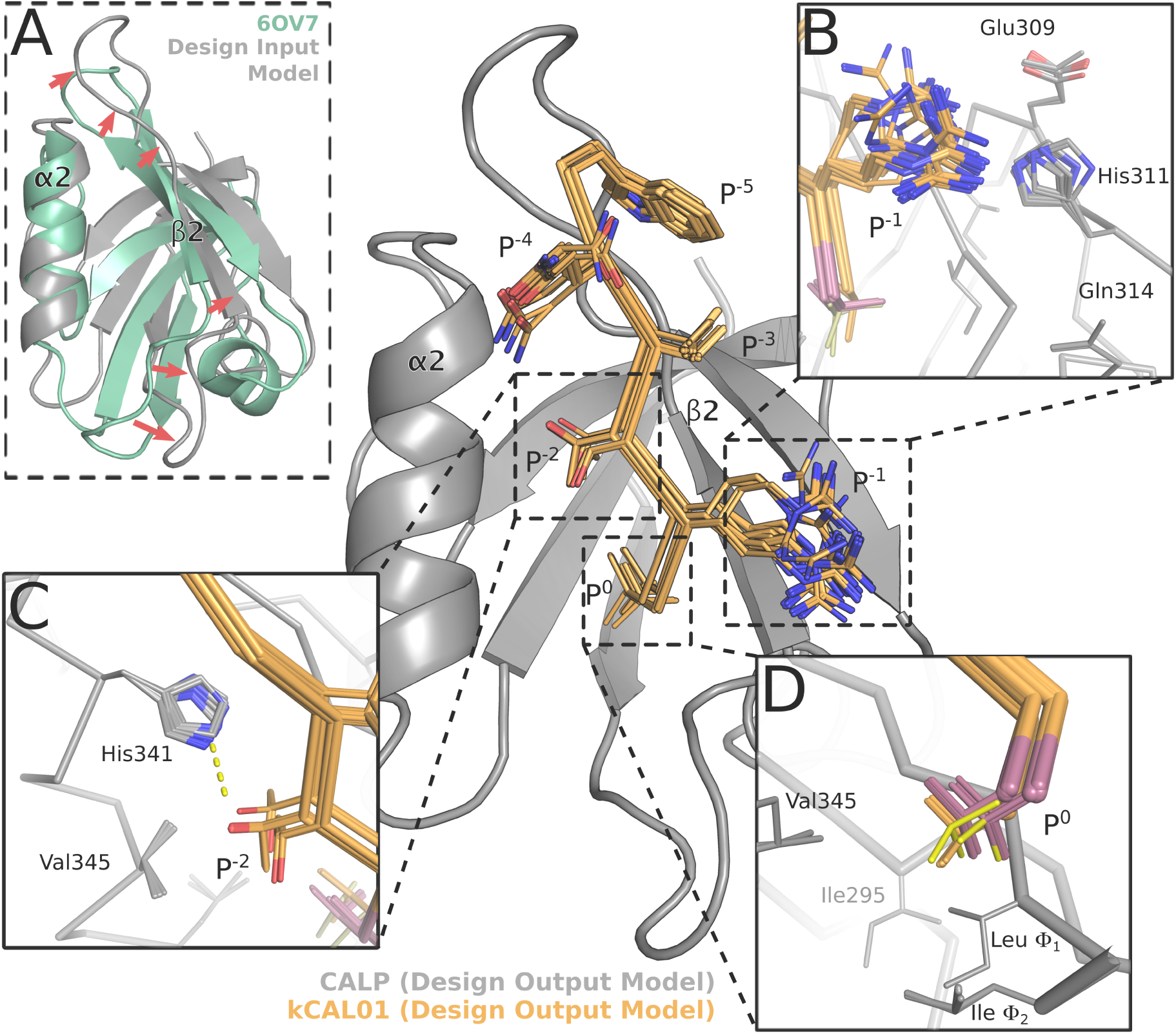
Structural analysis of the kCAL01 design models^2^ reveals similarities to the CALP:kCAL01 crystal structure. The design output model ensemble^2^ of CALP (gray) bound to kCAL01 (orange) closely resembles the bound CALP:kCAL01 crystal structure (Figure 1). (A) Comparison of the design input model (gray) and 6OV7 crystal structure (green) CALP conformations shows significant shifts (red arrows) in strand *β*2 and the *β*2-*β*3 loop. These shifts greatly expand the binding cleft between helix *α*2 and strand *β*2. The shift in *β*2-*β*3 loop conformation is a result of MD refinement, as it is not present in the structure of 2LOB before refinement (See Figure S2). The 100 conformations in the design output model ensemble capture interactions between (B)Arg P^-1^ and His311, as well as (C) the interaction between Thr P^-2^ and His341. (D) The design output ensemble models conformational heterogeneity at P^0^, suggesting that modeling of entropy at this site was important for this design’s success. The **t, p**, or **m** rotamers of P^0^ are shown in red, orange, and yellow, respectively.

We hypothesized that these differences were inherited from the CALP:CFTR design input model, generated during MD refinement^2,25^ of the bound NMR structure of CALP:CFTR (PDB ID: 2LOB).^25^ In order to test this hypothesis, we aligned both 2LOB and the design input model to 6OV7 by the main-chain atoms of secondary-structure elements *α*2 and *β*2 which revealed good alignment quality, with an RMSD of 0.69 and 0.94 Å, respectively. Overall, 2LOB is more relaxed than the bound CALP:kCAL01 structure, with slight outward shifts and changes in angle in both helix *α*2 and strand *β*2 that result in an apparent expansion of the hydrophobic pocket that interacts with peptide residue P^0^ (Figure S2). However, the loop connecting *β*2 and *β*3 shows good correspondence between the CALP:CFTR NMR structure and the CALP:kCAL01 crystal structure (Figure S2). The change in loop conformation and significant reorientation of strand *β*2 is a result of MD refinement, and does not appear in either the unrefined NMR structure (2LOB), the CALP:kCAL01 crystal structure (6OV7), or indeed in the CALP:iCAL36 structure (4E34). Therefore, we conclude that deviations in CALP conformation observed in the design output model of Ref.^2^ were inherited from MD refinement of a relaxed NMR structure.

Nonetheless, this NMR-based design model captured key structural and ensemble properties of the CALP:kCAL01 complex,^80^ which allowed OSPREY to design kCAL01, the most binding efficient inhibitor of CALP to date. Almost all members of the design ensemble predict favorable interactions between Arg P^-1^ and His311 (Figure 7B). Additionally, all three rotamers of Val P^0^ (**t, p**, and **m**) appear in the design output ensemble, indicating that the *K\** algorithm successfully identified sequences with multiple low-energy states (Figure 7D). A significant subset of the design ensemble captures the important hydrogen bond between His341 and Thr P^-2^, indicating that the design of kCAL01 captured key components of the class 1 PDZ binding geometry^28^ (Figure 7C). Interestingly, some members of the design output ensemble do not contain the hydrogen bond between Histidine 341 and Threonine P^-2^, consistent with observations of hydrogen bond breaking and reformation from experimental^81^ and MD simulation^82–85^ studies. Indeed, solution NMR studies of ubiquitin showed that threonine residues occupy multiple rotameric states.^86^ Overall, the crystal structure and design ensemble are quite quantitatively and qualitatively similar, despite differences in CALP structure, indicating that the design presented in Ref.^2^ succeeded in capturing important structural and dynamic interactions. We conclude that these features allowed OSPREY to design kCAL01, the most binding efficient inhibitor of CALP to date with rescue activity for F508del-CFTR, a disease-associated variant present in approximately 90% of CF patients. Key interactions and entropic effects predicted by the OSPREY design model are supported by the new crystal structure and landscape analysis presented herein.

### 3.4 PDZ domain energy landscapes

Modeling of energy landscapes complements traditional structural analysis of CALP:peptide crystal structures, and provides a novel way to probe the conformational distribution available to the protein complex. We submit that these tools may prove useful for analyzing PDZ:peptide complexes in general. To investigate differences in conformational distributions for PDZ domains, we computed partition functions and energy landscapes for 10 structures of bound PDZ:peptide complexes. A preliminary analysis of these landscapes reveals a general trend of loss of entropy upon binding for both PDZ and peptide ligands, similar to that observed for CALP:kCAL01 and CALP:iCAL36. Additionally, these supplementary energy landscapes predict no significant conformational heterogeneity for any studied system at the peptide position P^0^ in the bound state, contrasting with the heterogeneity we observed for CALP:kCAL01 at Val P^0^. This raises the intriguing possibility that, similarly to kCAL01 for CALP, more binding efficient inhibitors for other PDZ domains could be designed by maximizing relative entropy at P^0^. We include these energy landscapes for the scientific communtiy in SI Section S1.3 in the hope that these insights and data may be of further benefit.

## 4 Conclusion

In this work we investigated the basis for the binding efficiency of kCAL01, an OSPREY-designed peptide inhibitor of CALP that rescued functional CFTR activity as assessed by *in vitro* Ussing chamber assays.^2^ Based on structure and energy landscape analysis of the new crystal structure of CALP:kCAL01, we conclude that the comparative binding efficiency of kCAL01 stems from entropic effects at P^0^ and substitutions that result in more favorable energetic interactions at modulator residues. This conclusion is supported not only by comparative analysis of the CALP:kCAL01 and CALP:iCAL36 crystal structure conformations, but also by investigating energy landscapes for each ensemble model. We used energy landscape analysis enabled by the *MARK\** algorithm^4^ in OSPREY to provably approximate the energies of all conformations in each ensemble model, generated by assigning flexibility to residues in each crystal structure. These landscapes probed local residue conformational heterogeneity, and enabled us to approximate binding thermodynamics to correctly predict that kCAL01 binds more tightly to CALP than does iCAL36. We conclude that modeling of energy landscapes complemented traditional structural analysis of CALP:peptide crystal structures, and provided a novel way to probe the conformational distribution available to the protein complex. Modeling energy landscapes may prove useful for analyzing PDZ:peptide complexes in general, and hence we provide energy landscapes for 10 additional bound PDZ:peptide complexes. Finally, we show that our successful design of kCAL01^2^ was a result of effective modeling of both energetic and ensemble properties of CALP:peptide binding.

## Supporting Information Available

Supplementary figures and tables of structures (Figures S1, S2), previous biochemical characterization (Table S1), computational design flexibility (Table S2), and energy landscapes (Figures S3-S14).

## Acknowledgement

The authors thank Carrie Ann Davison and Dr. Mark Spaller for kCAL01 peptide synthesis; Terrence Oas, Jane and Dave Richardson, Nathan Guerin, Hong Niu, and all members of the lab for helpful discussion and comments; the NIH (R01-GM078031, R01-GM118543 to BRD; R01-DK101541, P20-GM113132, P30-DK117469, T32-GM008704 to DRM) for funding; and the Stanford Synchrotron Radiation Lightsource (SSRL) staff, especially Irimpan I. Mathews. Use of the Stanford Synchrotron Radiation Lightsource, SLAC National Accelerator Laboratory, is supported by the U.S. Department of Energy, Office of Science, Office of Basic Energy Sciences under Contract No. DE-AC02-76SF00515. The SSRL Structural Molecular Biology Program is supported by the DOE Office of Biological and Environmental Research, and by the National Institutes of Health, National Institute of General Medical Sciences.

