## Supplemental Information for "Computational Analysis of Energy Landscapes Reveals Dynamic Features that Contribute to Binding of Inhibitors to CFTR-Associated Ligand"

### S1 Supplementary Information

#### S1.1 Structural analysis

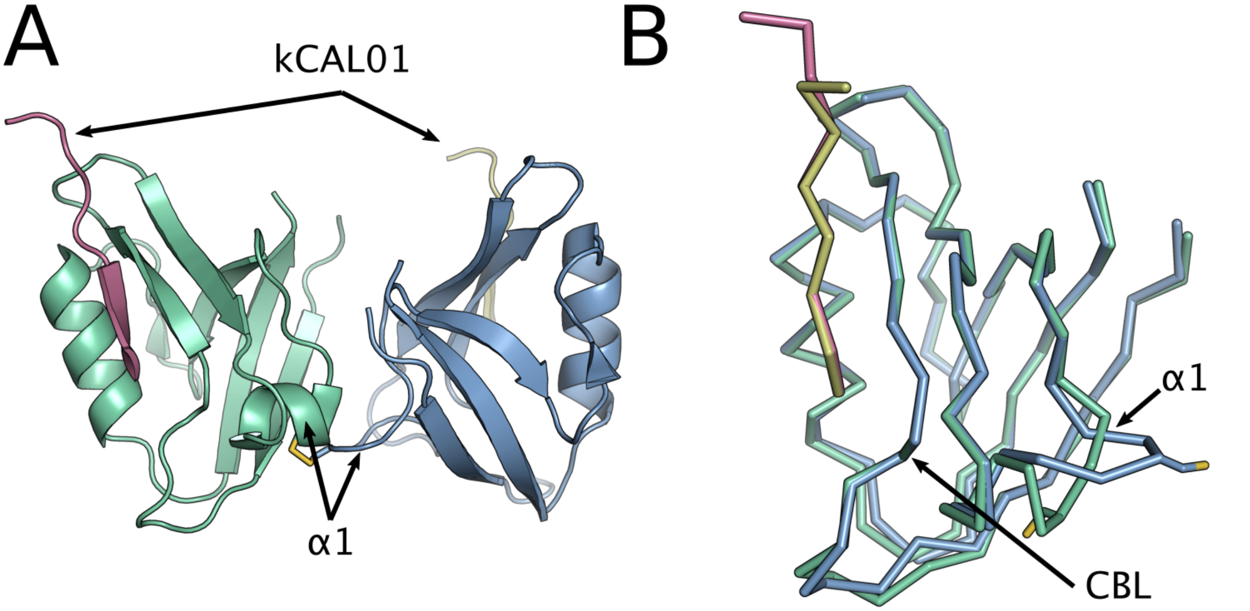


Figure S1: **Crystal structure of CALP:kCAL01 (PDB ID: 6OV7).** CALP and kCAL01 structure models shown in green and pink, respectively (protomer A), and blue and yellow, respectively (protomer B). (A) The asymmetric unit contains two bound CALP:kCAL01 protomers, shown in cartoon representation. Note the distortion of the protomer B helix $\alpha 1$ caused by an interprotomer cysteine bond. (B) Protomer structures aligned by main chain atoms, shown by $C_\alpha$ trace. Structures align well, although notable differences in CALP conformation are found at helix $\alpha 1$ and adjacent to the carboxylate binding loop (CBL).

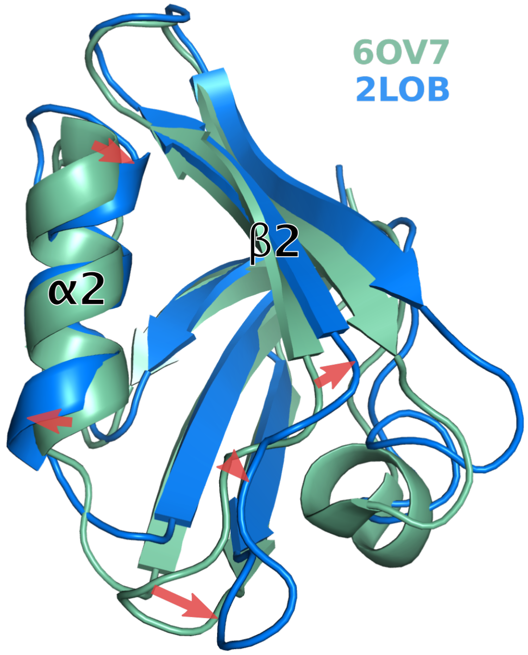


Figure S2: **Structural comparison of the CALP:kCAL01 (6OV7) crystal structure and CALP:CFTR (2LOB) NMR structure.** The CALP conformation in the NMR structure of CALP:CFTR[25] (blue) displays a more relaxed binding cleft than in the CALP:kCAL01 (6OV7) crystal structure (green). Comparison of these two structures shows moderate shifts (red arrows) in strand $\beta 2$ and helix $\alpha 2$. These shifts expand the binding cleft between helix $\alpha 2$ and sheet $\beta 2$.

**Table S1: Inhibition constants ($K_i$) and molecular weights for peptide inhibitors of CALP.** The name, inhibition constants, and molecular weights are shown for selected inhibitors of CALP. In addition to kCAL01 (shown here), 11 additional 6-mer peptides are reported in Ref.,[2] all of which have lower affinity for CALP than kCAL01 by at least 5-fold ($K_i$). Addition of a fluorescein dye moiety (F*) has been shown to improve the binding to CALP of most peptide inhibitors by approximately 10-fold.[26,30]

| Inhibitor name | Inhibitor Sequence | CALP $K_i$ ($\mu$M) | Molecular Weight (g/mol)[a] |
|---|---|---|---|
| $\text{CFTR}_{10}$[c] | TEEEVQDTRL | $420 \pm 80$ | 1218.6 |
| $\text{iCAL36}_6$[b,e] | WPTSII | $32.8 \pm 0.3$ | 818.4 |
| $\text{iCAL36}_{10}$[c] | ANSRWPTSII | $22.6 \pm 8.0$ | 1143.6 |
| F*-$\text{iCAL36}_{10}$[b,f] | F*-ANSRWPTSII | $1.3 \pm 0.1$ | 1501.9 |
| $\text{kCAL01}_6$[d] | Ac-WQVTRV | $2.3 \pm 0.2$ | 829.4 |

[a]Calculated with reduced cysteines. [b]Values previously reported in Ref.[26] [c]Values previously reported in Ref.[27] [d]Values previously reported in Ref.[2] [e]Peptides include an N-terminal cysteine to permit labeling. [f]$K_d$ values are shown for this entry.

Table S2: **Flexible residues for energy landscape computations.**

| PDB ID | Complex name (protein:ligand) | Protein flexible residues (⟨Chain⟩⟨#⟩) | Ligand flexible residues (⟨Chain⟩⟨#⟩) | Omitted residues (⟨Chain⟩⟨#⟩) |
|---|---|---|---|---|
| 6OV7 | CALP:kCAL01 | A296, A300, A301, A303, A306, A311, A314, A341, A345 | C5, C6, C7, C8, C9, C10 | A276, C2 |
| 4E34 | CALP:iCAL36 | A319, A309, A349, A322, A308, A353, A311, A314, A304 | C5, C6, C7, C8, C9, C10 | A284 |
| 1BE9 | PSD-95 PDZ3:CRIPT | A326, A328, A331, A339, A372, A376, A380 | B5, B6, B7, B8, B9 | A301, A302 |
| 1MFG | Erbin PDZ:ErbB2 | A1294, A1296, A1302, A1316, A1347, A1351 | B1250, B1251, B1252, B1253, B1254, B1255 | A1277, B1247 |
| 3NGH | PDZK1 PDZ1:SR-B1 | B23, B25, B26, B28, B68, B72, B76 | A106, A107, A108, A109, A110, A111 | B6, A103 |
| 3RL7 | hDLG1 PDZ1:APC | A237, A239, A243, A244, A256, A289, A293 | G2838, G2839, G2840, G2841, G2842, G2843 | A220, G2837 |
| 4K6Y | CALP:iCAL36-Q | B304, B308, B309, B311, B314, B319, B322, B349, B353 | D5, D6, D7, D8, D9, D10 | B284 |
| 4K72 | CALP:iCAL36-VQD | B304, B308, B309, B311, B314, B319, B322, B349, B353 | D5, D6, D7, D8, D9, D10 | B284 |
| 4K75 | CALP:iCAL36-QDTRL | A304, A308, A309, A311, A314, A319, A322, A349, A353 | B5, B6, B7, B8, B9, B10 | A284, B4 |
| 4K76 | CALP:iCAL36-TRL | D304, D308, D309, D311, D314, D319, D322, D349, D353 | H5, H6, H7, H8, H9, H10 | D284, D327 |
| 4WYU | Scribble PDZ3:peptide | A27, A29, A34, A49, A81, A85 | D−5, D−4, D−3, D−2, D−1, D0 | A3 |
| 5VWK | Scribble PDZ1:Beta-PIX | A741, A748, A749, A751, A761, A793, A797 | H158, H159, H160, H161, H162, H163 | A714, A716 |

#### S1.2 Bound landscapes for kCAL01 and iCAL36

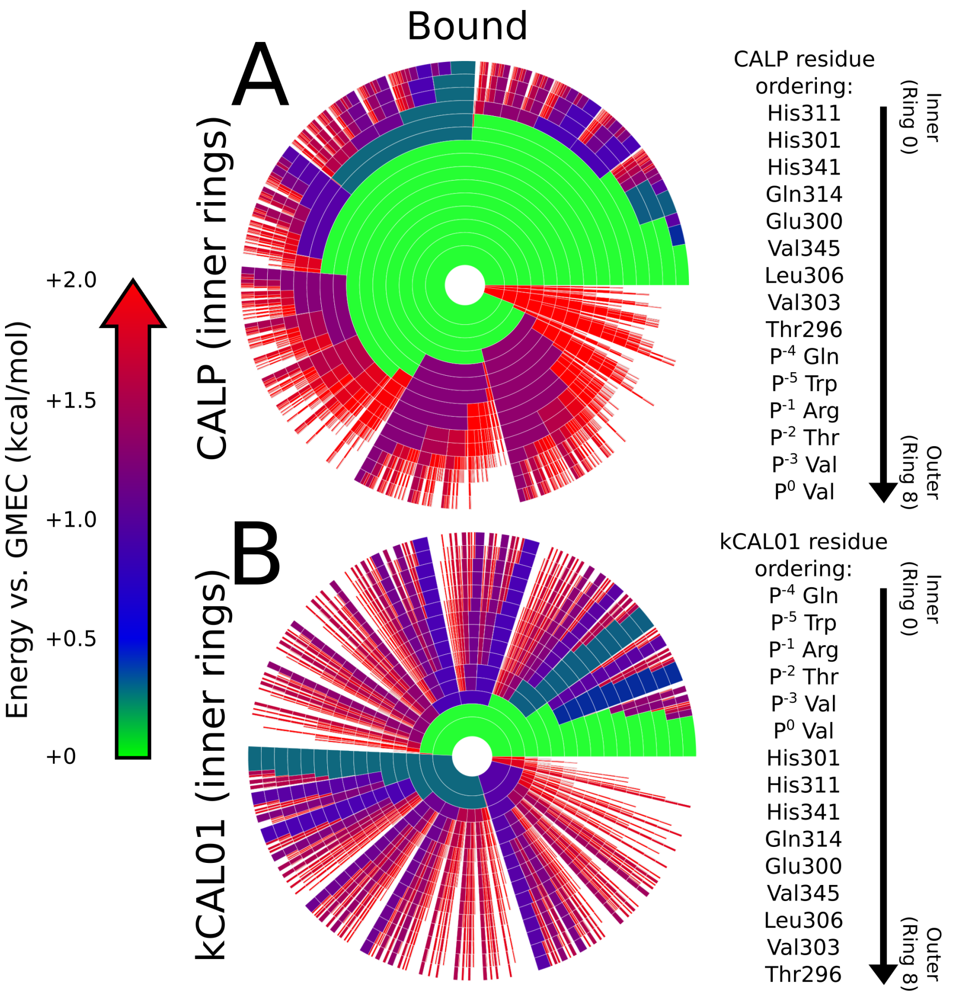


Figure S3: **Energy landscapes for the bound CALP:kCAL01 complex.** Upper bounds on the Boltzmann-weighted partition function computed using the *MARK** algorithm[4] for a 15-residue design at the protein-protein interface of CALP:kCAL01 are shown as colored ring charts. (A) Bound landscape (protein:inhibitor) with CALP residues shown in the inner rings, kCAL01 residues shown in the outer rings. (B) Bound landscape (protein:inhibitor) with kCAL01 residues shown in the inner rings, CALP residues shown in the outer rings.

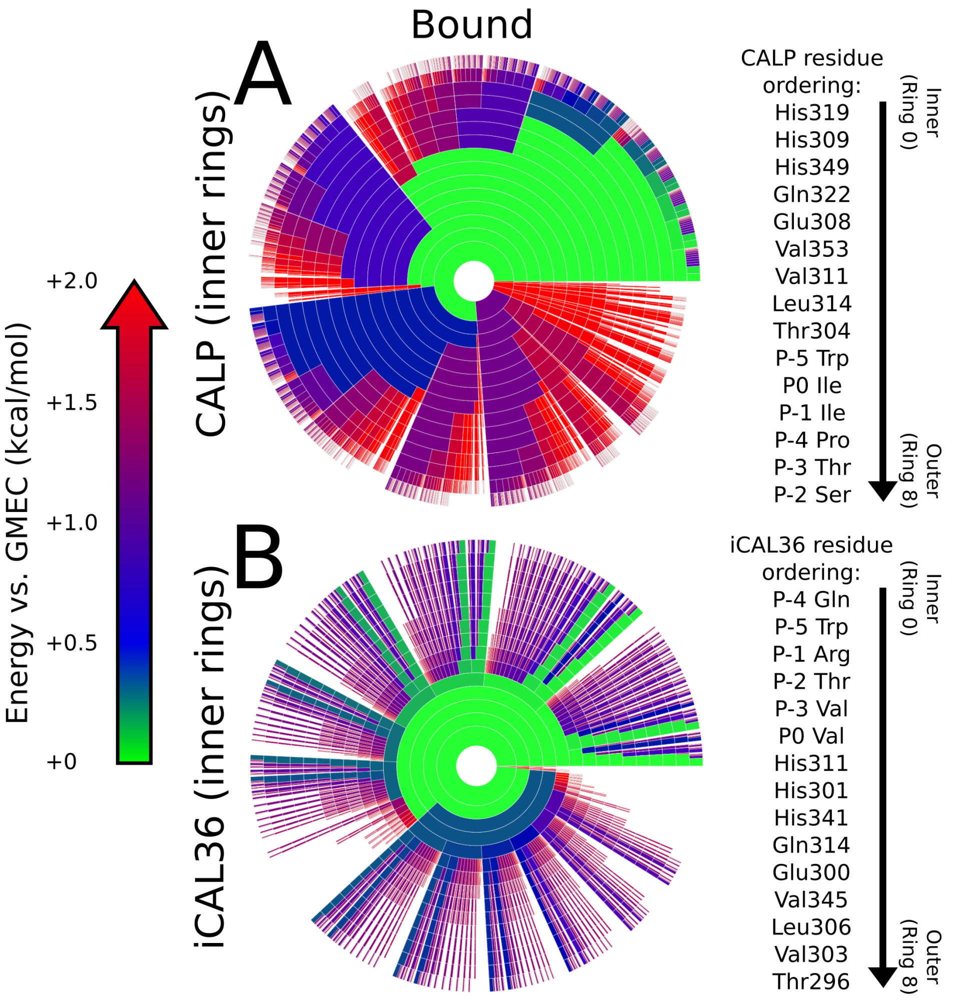


Figure S4: **Energy landscapes for the bound CALP:iCAL36 complex.** Upper bounds on the Boltzmann-weighted partition function computed using the *MARK** algorithm[4] for a 15-residue design at the protein-protein interface of CALP:iCAL36 are shown as colored ring charts. (A) Bound landscape (protein:inhibitor) with CALP residues shown in the inner rings, iCAL36 residues shown in the outer rings. (B) Bound landscape (protein:inhibitor) with iCAL36 residues shown in the inner rings, CALP residues shown in the outer rings.

#### S1.3 Energy landscapes for other PDZ domain models

Here we present energy landscapes of PDZ:peptide binding for various PDZ complexes. In particular, we investigate PDZ domains from the PSD-95, Erbin, PDZK1 (or NHERF3), Discs large homolog 1 (DLG1), Scribble, and CAL proteins. For each system, landscapes were generated as described in Sections 2.2 and 2.3 of the main manuscript. Briefly, the bound and unbound states were defined using one protein complex from the asymmetric unit of each crystal structure. Hydrogens were added using MolProbity,[67] and selected residues (See Figure S2) were designated as continuously flexible using continuous rotamers.[68,69] Partition functions were approximated to a guaranteed accuracy of $\varepsilon < 0.1$, using *MARK*$^*$ in OSPREY, and energy landscapes were generated using the same protocol as those in the main manuscript.

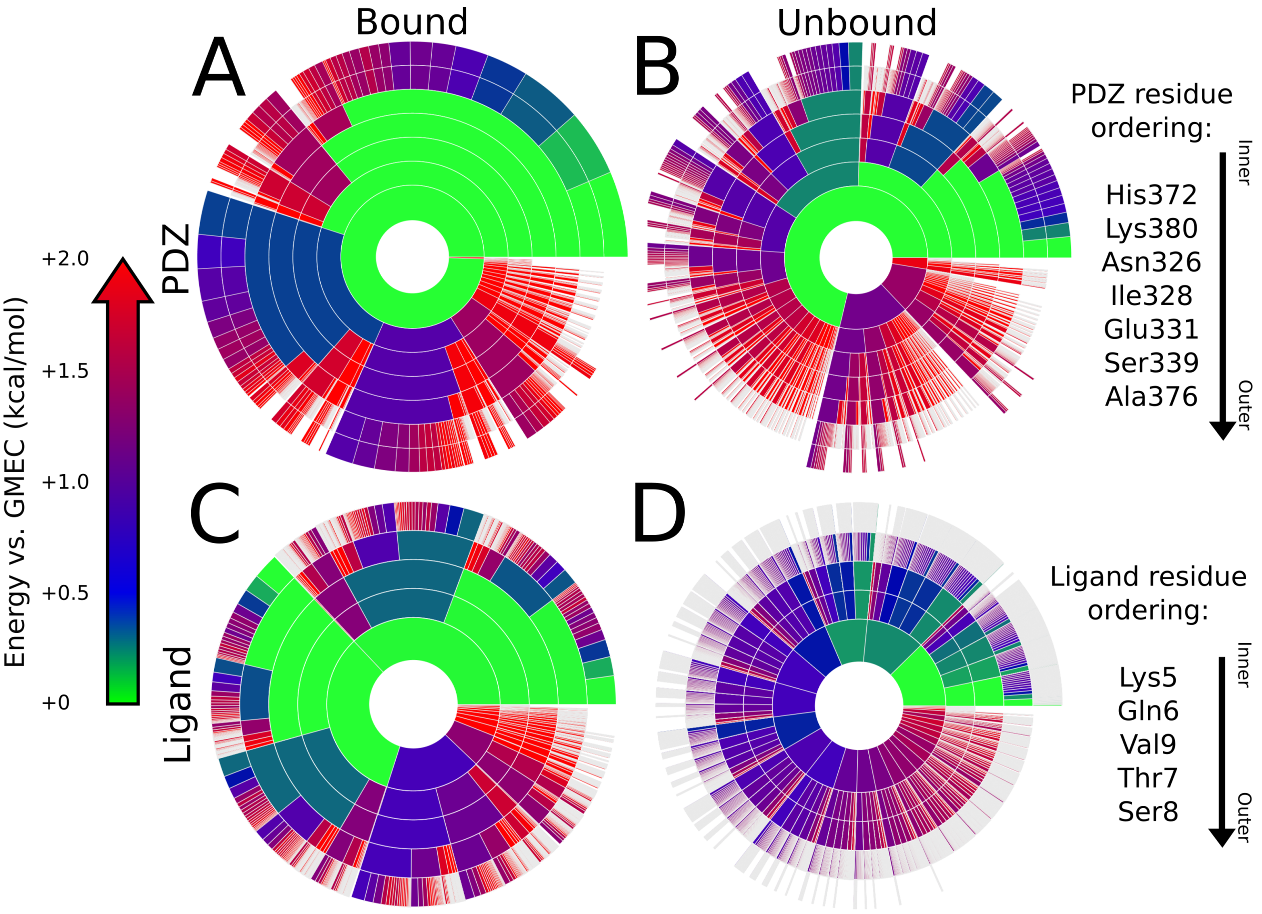


Figure S5: **Energy landscapes of binding for PSD-95 PDZ3:CRIPT (PDB: 1BE9) binding interface.** Upper bounds on the Boltzmann-weighted partition function computed using the *MARK** algorithm[4] in OSPREY[3] for a design at the protein-protein interface of PSD-95 PDZ3:CRIPT (PDB: 1BE9) shown as colored ring charts. A brief explanation of the ring chart diagram can be found in Section 2.4. Energy landscapes for the PDZ binding domain in the bound (A) and unbound (B) states, along with landscapes for the peptide ligand in the bound (C) and unbound (D) states are shown.

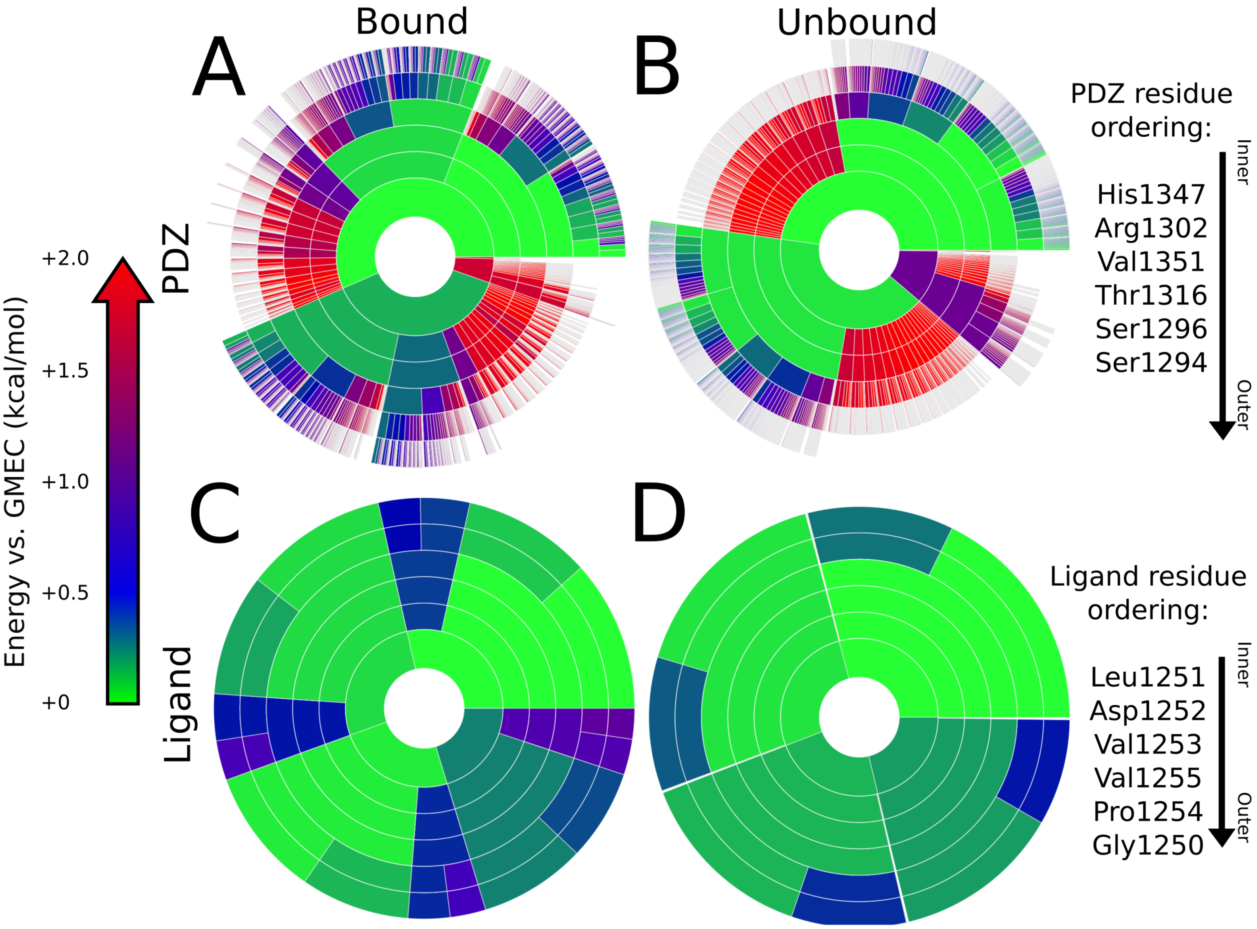


Figure S6: **Energy landscapes of binding for Erbin PDZ:ErbB2 (PDB: 1MFG) binding interface.** Upper bounds on the Boltzmann-weighted partition function computed using the *MARK** algorithm[4] in OSPREY[3] for a design at the protein-protein interface of Erbin PDZ:ErbB2 (PDB: 1MFG) shown as colored ring charts. A brief explanation of the ring chart diagram can be found in Section 2.4. Energy landscapes for the PDZ binding domain in the bound (A) and unbound (B) states, along with landscapes for the peptide ligand in the bound (C) and unbound (D) states are shown.

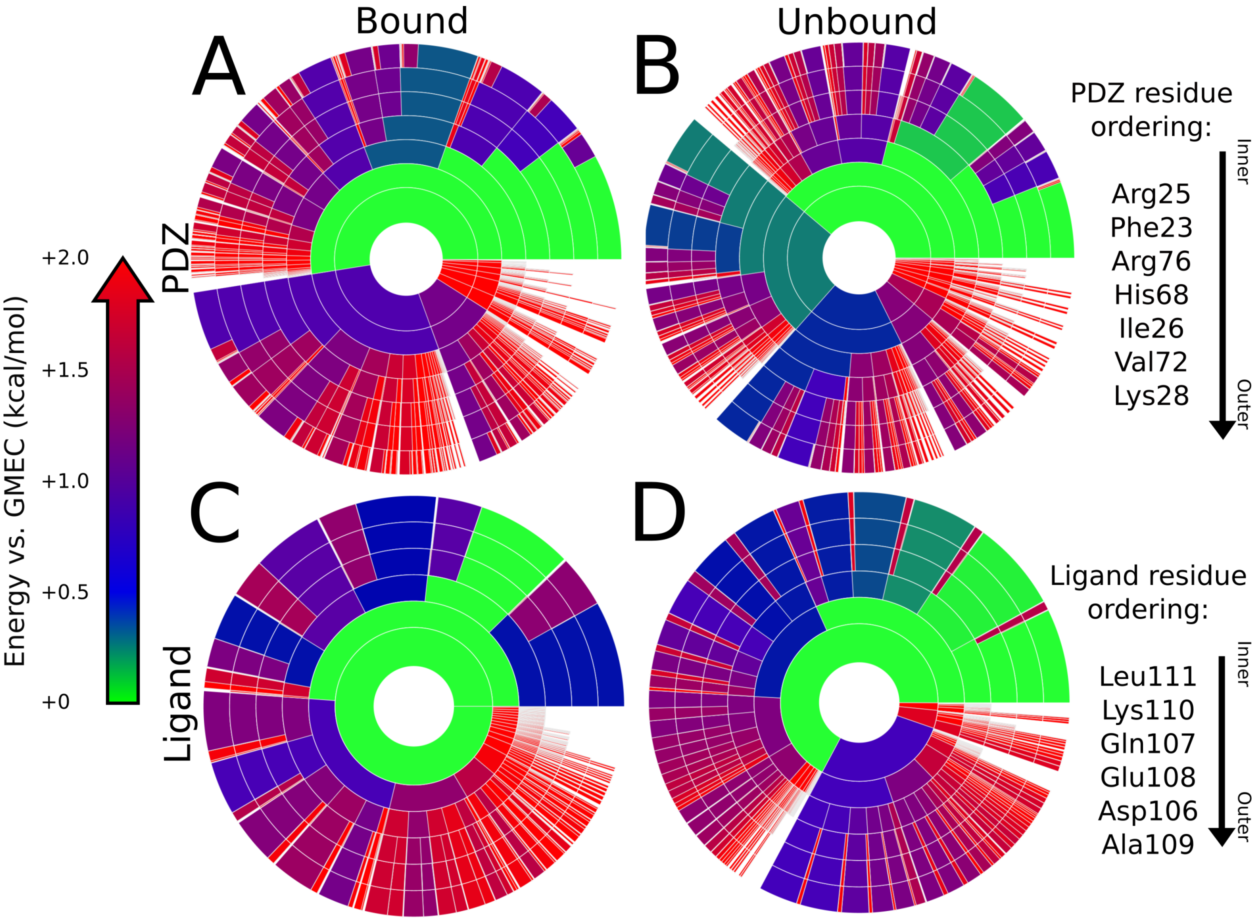


Figure S7: **Energy landscapes of binding for PDZK1 PDZ1:SR-B1 (PDB: 3NGH) binding interface.** Upper bounds on the Boltzmann-weighted partition function computed using the *MARK** algorithm[4] in OSPREY[3] for a design at the protein-protein interface of PDZK1 PDZ1:SR-B1 (PDB: 3NGH) shown as colored ring charts. A brief explanation of the ring chart diagram can be found in Section 2.4. Energy landscapes for the PDZ binding domain in the bound (A) and unbound (B) states, along with landscapes for the peptide ligand in the bound (C) and unbound (D) states are shown.

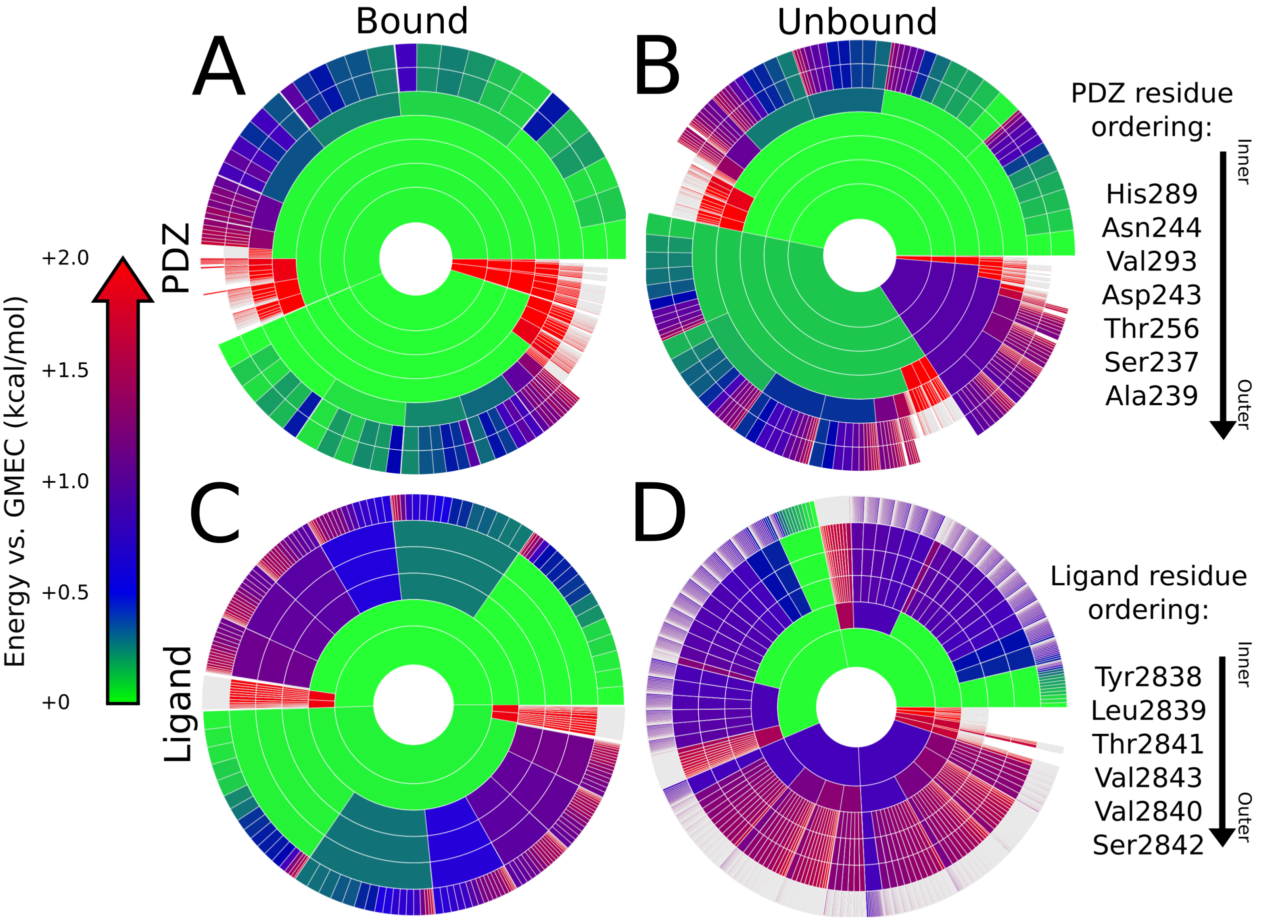


Figure S8: **Energy landscapes of binding for hDLG1 PDZ1:APC (PDB: 3RL7) binding interface.** Upper bounds on the Boltzmann-weighted partition function computed using the *MARK** algorithm[4] in OSPREY[3] for a design at the protein-protein interface of hDLG1 PDZ1:APC (PDB: 3RL7) shown as colored ring charts. A brief explanation of the ring chart diagram can be found in Section 2.4. Energy landscapes for the PDZ binding domain in the bound (A) and unbound (B) states, along with landscapes for the peptide ligand in the bound (C) and unbound (D) states are shown.

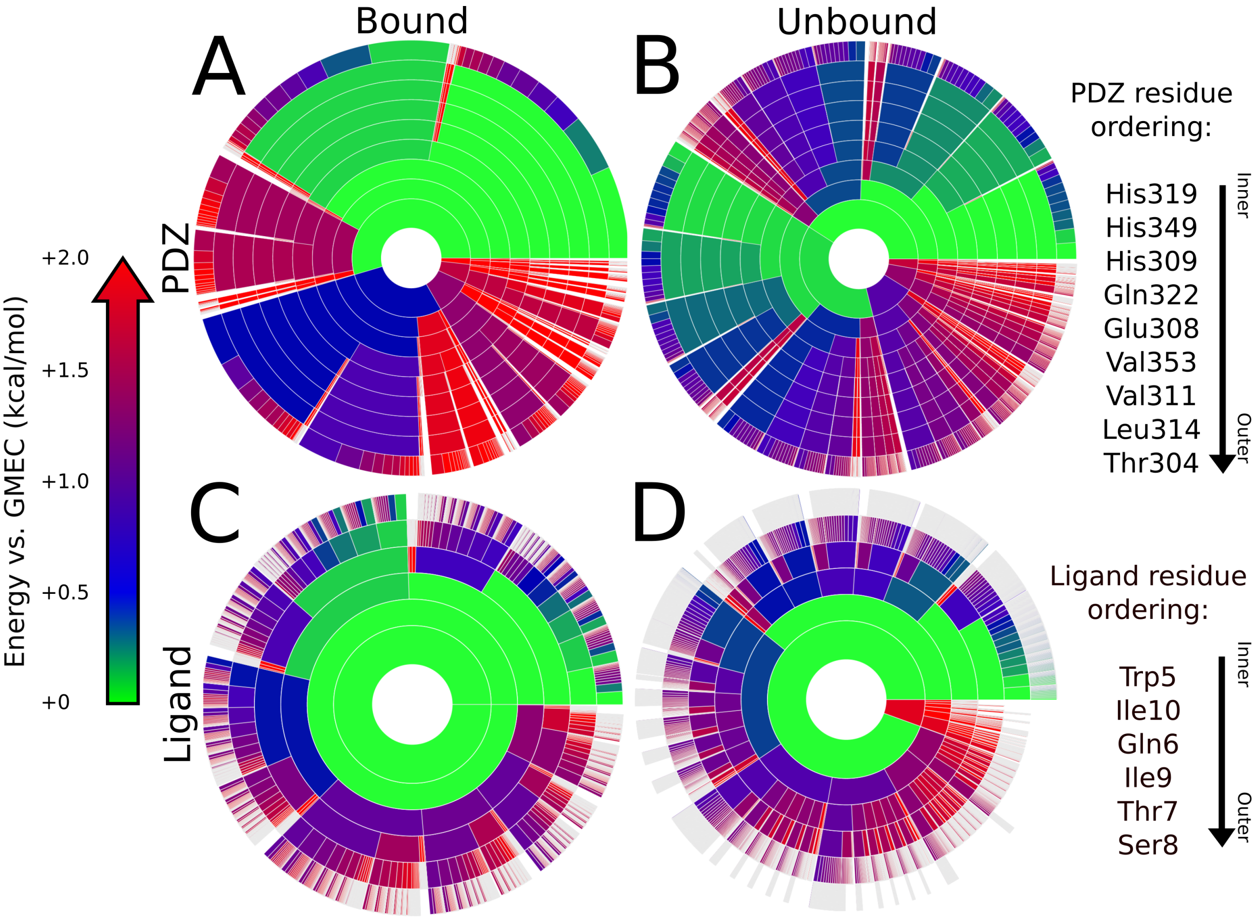


Figure S9: **Energy landscapes of binding for CALP:iCAL36-Q (PDB: 4K6Y) binding interface.** Upper bounds on the Boltzmann-weighted partition function computed using the *MARK** algorithm[4] in OSPREY[3] for a design at the protein-protein interface of CALP:iCAL36-Q (PDB: 4K6Y) shown as colored ring charts. A brief explanation of the ring chart diagram can be found in Section 2.4. Energy landscapes for the PDZ binding domain in the bound (A) and unbound (B) states, along with landscapes for the peptide ligand in the bound (C) and unbound (D) states are shown.

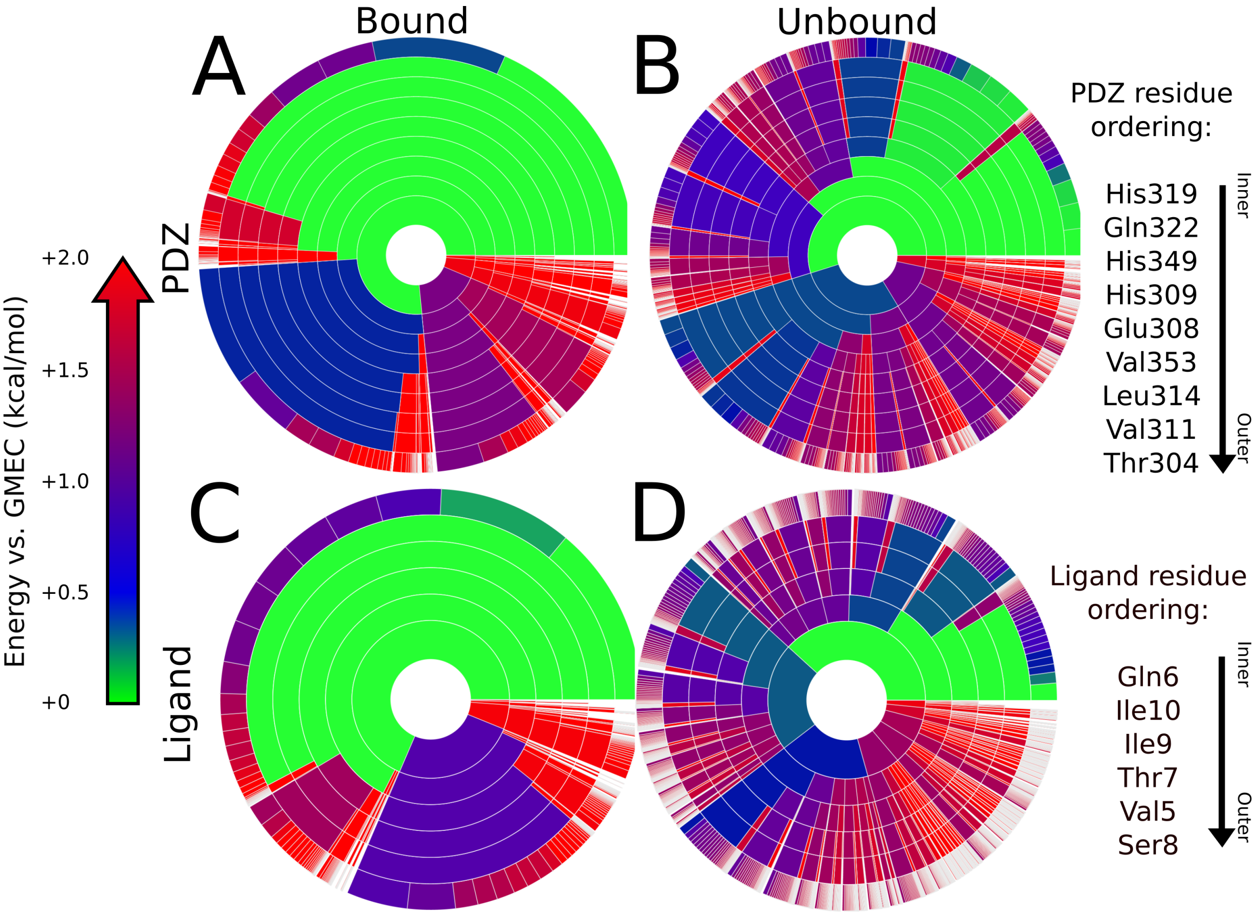


Figure S10: **Energy landscapes of binding for CALP:iCAL36-VQD (PDB: 4K72) binding interface.** Upper bounds on the Boltzmann-weighted partition function computed using the *MARK** algorithm[4] in OSPREY[3] for a design at the protein-protein interface of CALP:iCAL36-VQD (PDB: 4K72) shown as colored ring charts. A brief explanation of the ring chart diagram can be found in Section 2.4. Energy landscapes for the PDZ binding domain in the bound (A) and unbound (B) states, along with landscapes for the peptide ligand in the bound (C) and unbound (D) states are shown.

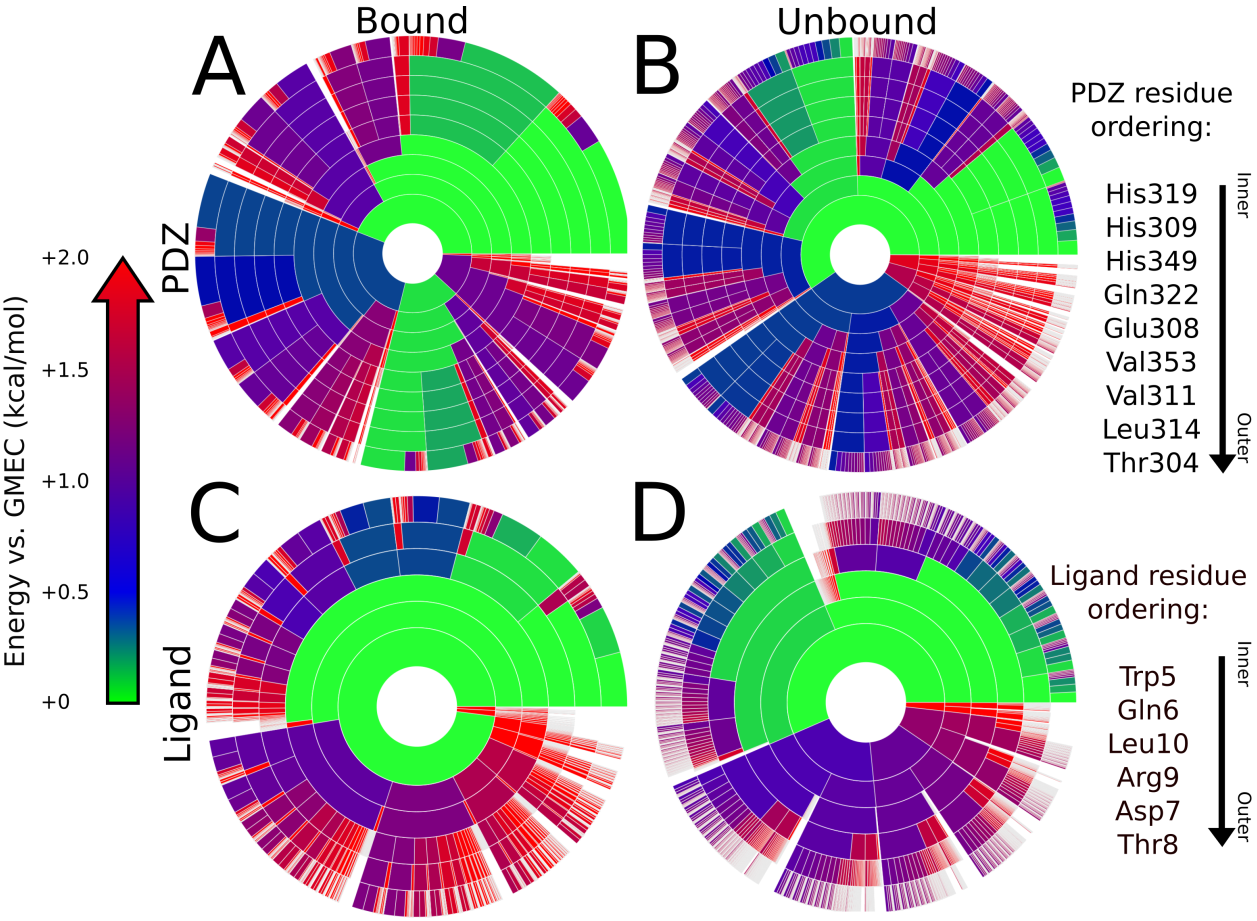


Figure S11: **Energy landscapes of binding for CALP:iCAL36-QDTRL (PDB: 4K75) binding interface.** Upper bounds on the Boltzmann-weighted partition function computed using the *MARK** algorithm[4] in OSPREY[3] for a design at the protein-protein interface of CALP:iCAL36-QDTRL (PDB: 4K75) shown as colored ring charts. A brief explanation of the ring chart diagram can be found in Section 2.4. Energy landscapes for the PDZ binding domain in the bound (A) and unbound (B) states, along with landscapes for the peptide ligand in the bound (C) and unbound (D) states are shown.

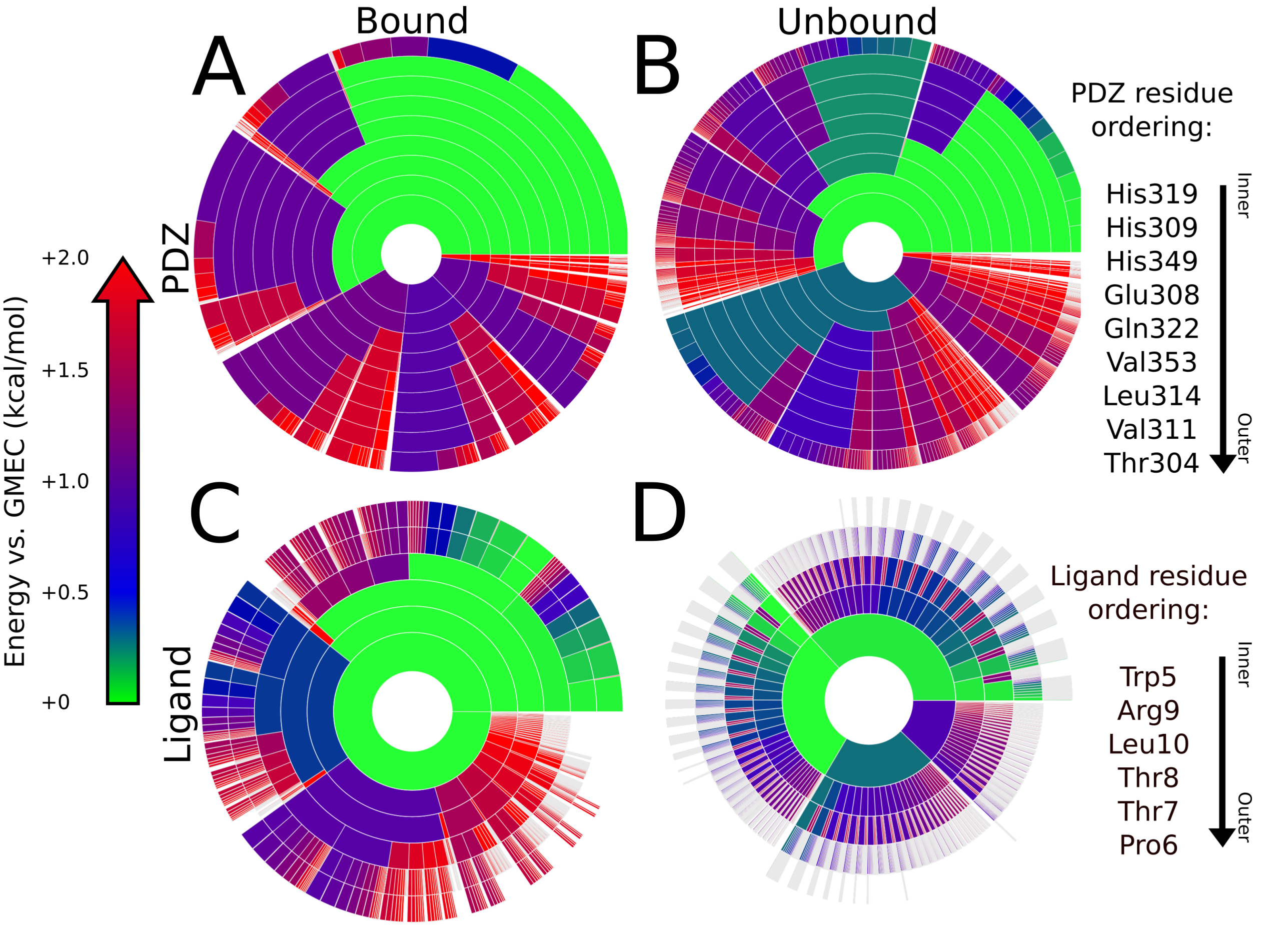


Figure S12: **Energy landscapes of binding for CALP:iCAL36-TRL (PDB: 4K76) binding interface.** Upper bounds on the Boltzmann-weighted partition function computed using the *MARK** algorithm[4] in OSPREY[3] for a design at the protein-protein interface of CALP:iCAL36-TRL (PDB: 4K76) shown as colored ring charts. A brief explanation of the ring chart diagram can be found in Section 2.4. Energy landscapes for the PDZ binding domain in the bound (A) and unbound (B) states, along with landscapes for the peptide ligand in the bound (C) and unbound (D) states are shown.

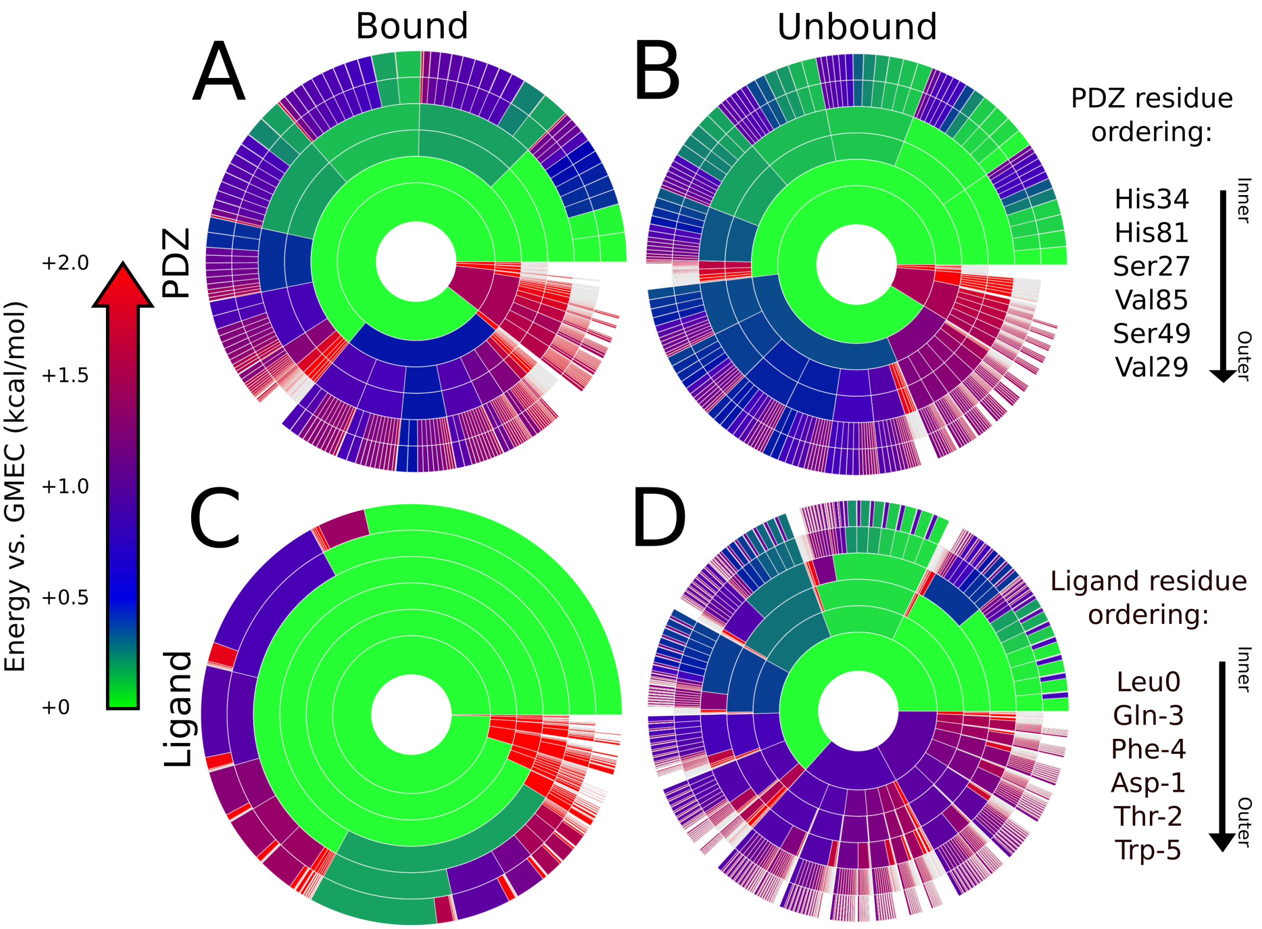


Figure S13: **Energy landscapes of binding for Scribble PDZ3:peptide (PDB: 4WYU) binding interface.** Upper bounds on the Boltzmann-weighted partition function computed using the *MARK** algorithm[4] in OSPREY[3] for a design at the protein-protein interface of Scribble PDZ3:peptide (PDB: 4WYU) shown as colored ring charts. A brief explanation of the ring chart diagram can be found in Section 2.4. Energy landscapes for the PDZ binding domain in the bound (A) and unbound (B) states, along with landscapes for the peptide ligand in the bound (C) and unbound (D) states are shown.

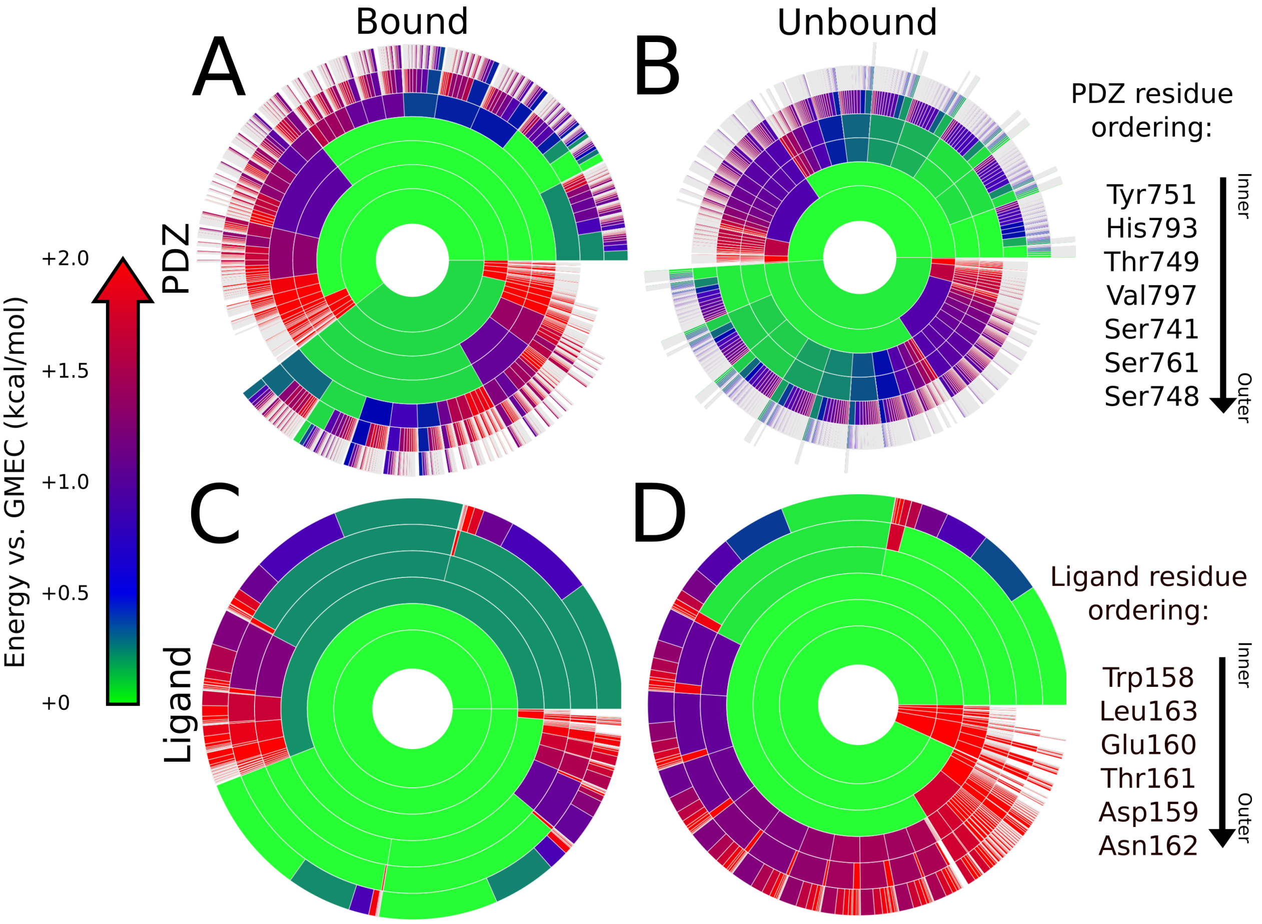


Figure S14: **Energy landscapes of binding for Scribble PDZ1:Beta-PIX (PDB: 5VWK) binding interface.** Upper bounds on the Boltzmann-weighted partition function computed using the *MARK** algorithm[4] in OSPREY[3] for a design at the protein-protein interface of Scribble PDZ1:Beta-PIX (PDB: 5VWK) shown as colored ring charts. A brief explanation of the ring chart diagram can be found in Section 2.4. Energy landscapes for the PDZ binding domain in the bound (A) and unbound (B) states, along with landscapes for the peptide ligand in the bound (C) and unbound (D) states are shown.
